# *Pseudomonas aeruginosa* mechanosensing controls cell polarity during twitching by activating two antagonistic response regulators

**DOI:** 10.1101/2022.11.15.516588

**Authors:** Marco J. Kühn, Henriette Macmillan, Lorenzo Talà, Yuki Inclan, Ramiro Patino, Xavier Pierrat, Zainebe Al-Mayyah, Joanne N. Engel, Alexandre Persat

## Abstract

The opportunistic pathogen *Pseudomonas aeruginosa* adapts to solid surfaces to enhance virulence and infect its host. Type IV pili (T4P), long and thin filaments that power surface-specific twitching motility, allow single cells to mechanosense surfaces. For example, cells sense T4P attachment to control the direction of twitching motility. In this process, they establish a local positive feedback that polarizes T4P distribution to the sensing pole. A complex chemotaxis-like system called Chp mediates this response. The signalling mechanism allowing for transduction of this spatially-resolved signal is however unresolved. Here we demonstrate that the two Chp response regulators PilG and PilH enable dynamic cell polarization by coupling their antagonistic functions on T4P extension. By precisely quantifying the localization of fluorescent protein fusions, we show that PilG polarizes in response to mechanosensing through phosphorylation by the histidine kinase ChpA. We find that PilH is not inherently required for reversals. However, PilH activation is necessary to break the local positive feedback established by PilG so that forward-twitching cells can reverse. To spatially resolve mechanical signals, Chp thus locally transduces signals with a main output response regulator, PilG. To respond to signal changes, Chp uses its second regulator PilH to break the local feedback. By identifying the molecular functions of two response regulators that dynamically control cell polarization, our work provides a rationale for the diversity of architectures often found in non-canonical chemotaxis systems.

## Introduction

Bacteria use mechanosensing to rapidly adapt to life on surfaces (1,2). Mechanosensory systems regulate virulence, adhesion, biofilm formation and surface-specific motility (3–6). The opportunistic pathogen *Pseudomonas aeruginosa* senses surfaces using extracellular filaments called type IV pili (T4P) (7,8). *P. aeruginosa* responds to T4P-mediated mechanosensing on two timescales. Mechanosensing controls the direction of surface-specific twitching motility within seconds in a process called mechanotaxis (9). It also regulates the transcription of a series of genes responsible for acute virulence and surface adaptation through the production of the second messenger cyclic adenosine monophosphate (cAMP) over minutes to hours (6,8,10,11).

To power surface-specific twitching motility, T4P successively extend, attach and retract, thereby pulling a cell forward (12,13). During twitching, individual *P. aeruginosa* cells can move forward persistently and reverse spontaneously or upon collision (9). To twitch forward, *P. aeruginosa* localizes the T4P extension motor PilB to the cell pole facing the direction of motion. We refer to this asymmetric protein and T4P localization as polarization (Figure 1A).

**Figure 1:**
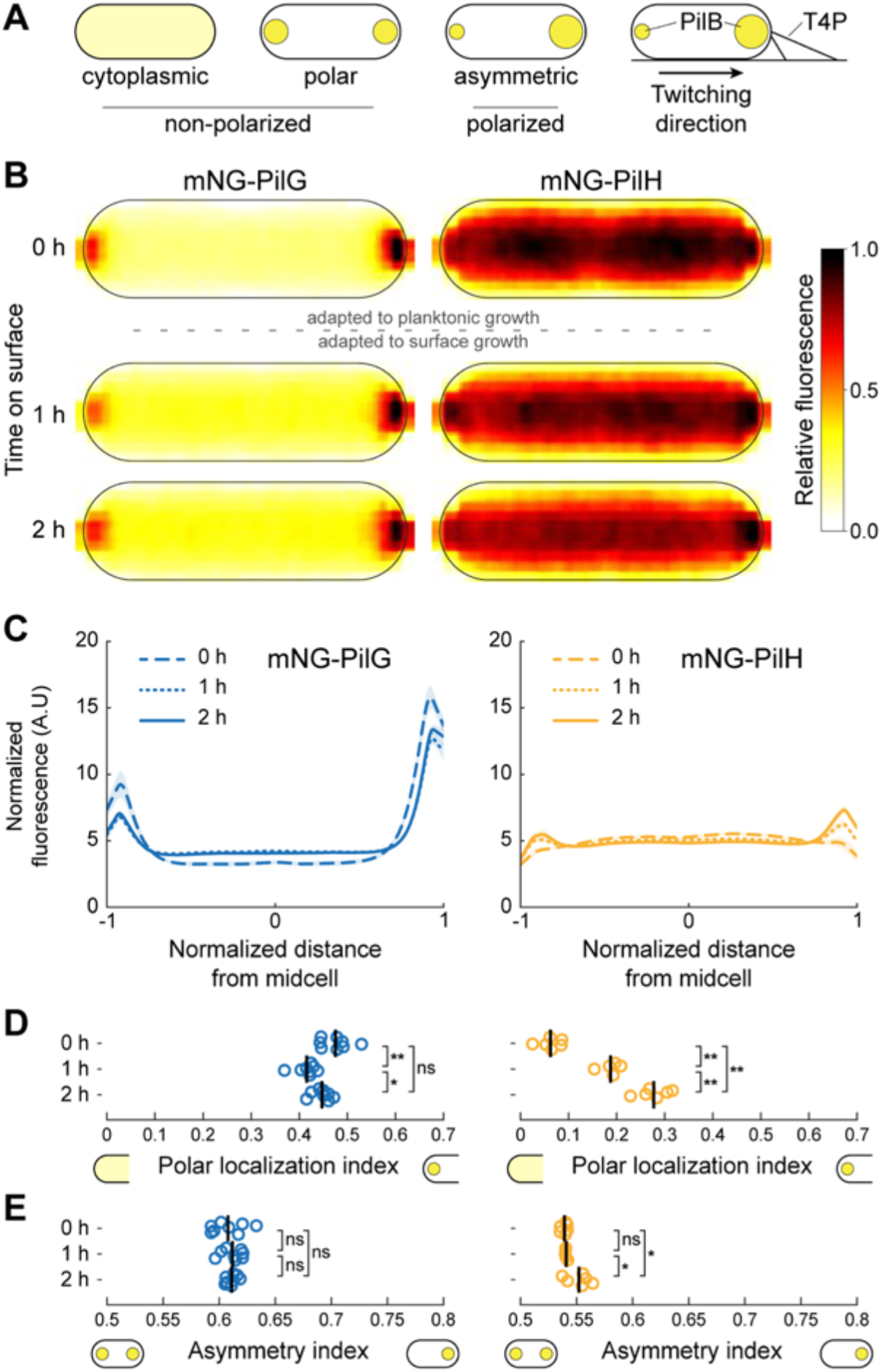
PilG and PilH localization during surface adaptation. (A) Schematic classification of localization patterns. The spot size indicates the relative protein concentration. (B) Average normalized fluorescent signal over 42 cells (bright pole to the right) for mNG-PilG and mNG-PilH transferred from liquid culture to a solid substrate and imaged immediately (0h) and after 1h and 2h. (C) Normalized average fluorescence profiles for mNG-PilG and mNG-PilH. The length of each cell is normalized so that the dim pole is positioned at *x* = −1, the bright pole at *x* = 1. The fluorescent profile for each cell is normalized to its total fluorescence. Solid lines: mean normalized fluorescence profiles across replicates. Shaded area, standard deviation across replicates. (D) The polar localization index measures the relative fraction of the fluorescence signal at the poles. PilG always localizes predominantly polarly. PilH is mostly cytoplasmic at 0h but becomes more polar over time. (E) Quantification of polarization. The asymmetry index measures the difference in fluorescent intensity between poles. In D and E, each circle corresponds to the population median of one biological replicate, and the vertical bars to the mean across biological replicates. *, p<0.05; **, p≤0.001; ns, not significant. For corresponding example images see Supplementary Figure 1A. For corresponding fluorescence measurements see Supplementary Figure 2A.

During mechanotaxis, T4P generate mechanical signal input and additional T4P deploy as output response, establishing a positive feedback loop. A chemotaxis-like system called Chp relays T4P signals to cellular components that ultimately control twitching and cAMP levels (14,15). In response to signal input at one pole, Chp controls the distribution of extension motor proteins to assemble T4P at that same pole. As a result of this polarized T4P deployment, single cells twitch persistently forward (9). To change twitching direction spontaneously or upon collision, cells reverse polarization (cf. Supplementary movie 1). To enforce the connection between T4P input and T4P polarization output, Chp employs two response regulators PilG and PilH that establish a local excitation - global inhibition signaling landscape (9). This signaling architecture is shared with higher order organisms: neutrophils, amoebae and yeast polarize via local excitation global inhibition during chemotaxis or cell division (16).

While the sensory system controlling flagellar rotation in *Escherichia coli*, the Che system, serves as a model chemotaxis system to understand more complex ones, it is more of an exception than the norm. Bacteria possess a wide diversity of chemotaxis systems with different inputs, outputs and overarching signaling mechanisms. Our knowledge of the dynamic control of these diverse chemotaxis-like systems is however limited. Chp shows important functional and structural differences with the canonical Che sensory system despite strong homology (17). First, Chp responds to a mechanical signal input from T4P but not to chemical ligands. Second, the Chp complex has more components than Che. CheA, the histidine kinases of the Che system, phosphorylates a single response regulator to relay input signals to its functional target. The histidine kinase ChpA comprises six phosphotransfer domains against just one for CheA (15). In addition, Chp possesses not one but two response regulators, PilG and PilH, with opposing functions in mechanotaxis and cAMP-dependent transcription (9–11,14). PilG promotes persistent forward twitching and cAMP production while PilH promotes reversals and reduces cAMP levels. However, how ChpA activates PilG and PilH to control cell polarization upon mechanosensing remains unresolved.

To guide twitching direction, T4P bind to the substrate and activate the Chp system, possibly through the interaction of the major pilin PilA and the Chp system receptor PilJ (8). As a result, PilJ stimulates ChpA autophosphorylation. ChpA then activates PilG and PilH presumably by phosphorylation (18). PilG and PilH activate their targets, which ultimately mediate polarization of the extension motor PilB and its activator FimX (9), although direct interactions have yet to be rigorously identified (19). PilG promotes recruitment of both PilB and FimX at the pole, while PilH inhibits it (9). As a consequence, T4P extend preferentially at that same pole, resulting in persistent forward twitching. Without a counterpart to PilG, i.e. in *pilH* deletion mutants, cells continue twitching in one direction and never reverse (9).

Despite homologies and analogies with other well-studied chemotaxis systems, how antagonistic response regulators established a stable, yet dynamic signaling landscape remains unknown. In particular, how PilG and PilH help transduce mechanical input into a polarization output that dynamically controls the direction of *P. aeruginosa* twitching is unresolved. Here, we investigate how phosphorylation by ChpA activates PilG and PilH to regulate cell polarity in response to mechanical signal input. We combined bacterial genetics with microscopy to determine the subcellular localizations and polarization of the two response regulators in their different active and inactive states. We suggest a model in which one regulator acts as primary signal relay while the other regulator only modulates the function of the first one.

## Results

### PilG and PilH localization during surface adaptation

To investigate how Chp activates response regulators upon surface contact, we monitored localization of functional mNeonGreen-tagged PilG and PilH as cells transition from liquid to a surface. Immediately after surface contact which reflects the state of planktonic cells that have yet to mechanosense, mNG-PilG predominantly localizes to the poles asymmetrically, *i.e*. PilG is polarized (Figure 1BC). We quantified subcellular localization by defining a polar localization index. PilG polar localization is largely insensitive to time on the surface with only a very small decrease (Figure 1D). To quantify protein polarization (Figure 1A), we defined an asymmetry index quantifying the intensity difference between poles. PilG is markedly and stably polarized, as the asymmetry index remains unchanged for at least 2h on surfaces (Figure 1E).

Unlike PilG, PilH is almost exclusively found in the cytoplasm early after surface contact (Figure 1B). Only a few cells display relatively dim mNG-PilH polar foci (Supplementary Figure 1A). mNG-PilH becomes increasingly localized to the poles after 1 and 2h of surface contact (Figure 1B). While PilH polar localization is always lower compared to PilG, the localization index increases over time on surface (Figure 1D). PilH asymmetry index is steadily close to 0.5, *i.e*. nearly symmetric, showing PilH does not polarize (Figure 1E). In summary, PilG remains strongly polarized, only switching polarization during reversals (9). PilH is largely cytoplasmic, but polar localization increases over time on surfaces without polarizing. We hypothesize that upon mechanosensing, ChpA causes polar recruitment of PilG and PilH.

### ChpA induces PilG polarization through phosphorylation

PilG preferentially localizes to the leading pole of twitching cells where T4P actively extend and retract (9). We therefore hypothesize that in response to mechanosensing with T4P, the histidine kinase ChpA polarizes PilG by phosphorylation. To characterize the role of phosphorylation in PilG polarization, we compared the localization of mNG-PilG between a *chpA* deletion mutant and wild type (WT). In *ΔchpA*, PilG polar localization decreases but is not entirely abolished (Figure 2AB). The asymmetry index is also lower in *ΔchpA* compared to WT, showing ChpA promotes PilG polarization (Figure 2C). This is consistent with a model where ChpA promotes PilG polarization towards the leading pole in response to T4P input (9).

**Figure 2:**
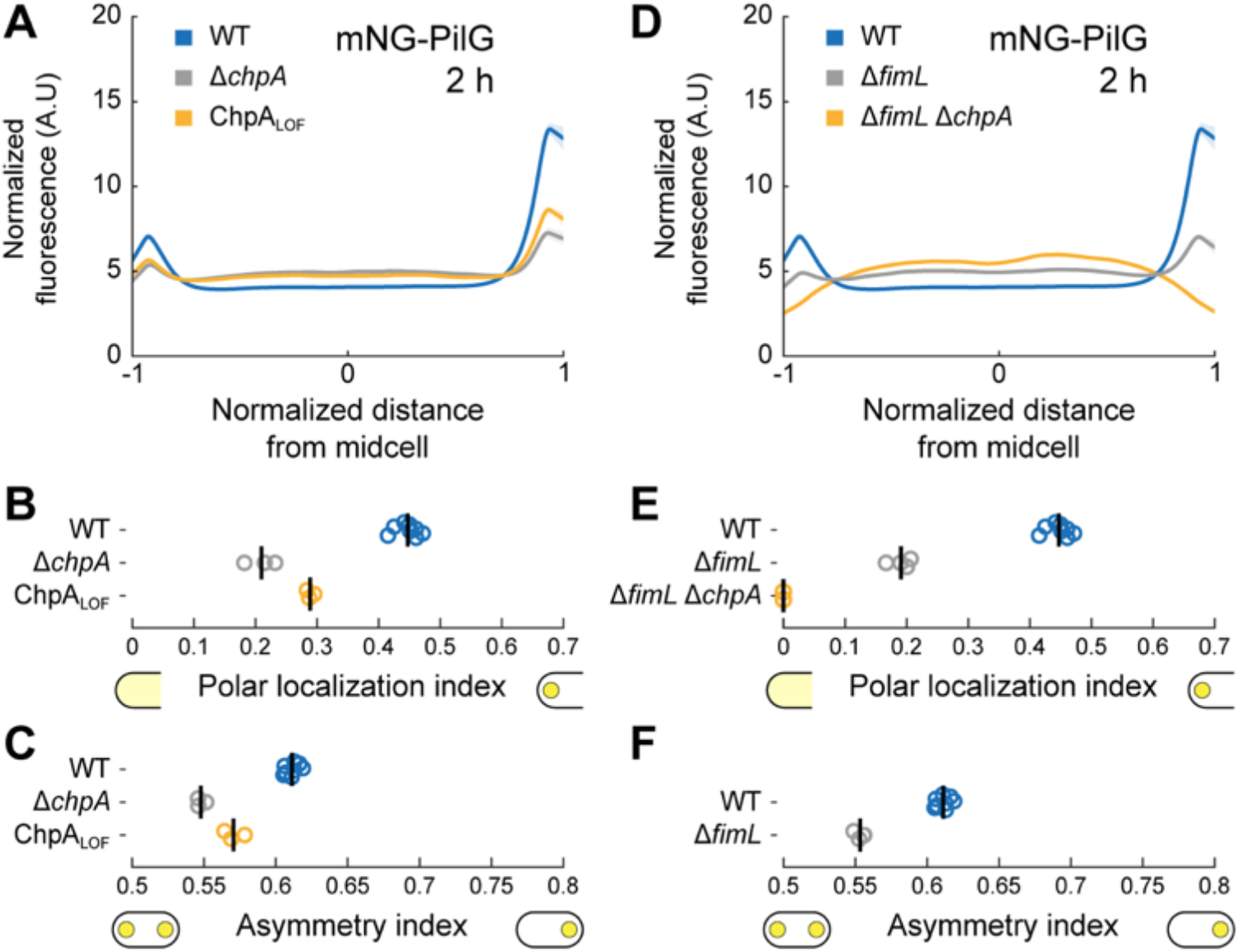
ChpA controls PilG polarization by phosphorylation. (A) Polar localization profiles of mNG-PilG in *chpA* mutants after 2h on surfaces. Solid lines, mean normalized fluorescence profiles across biological replicates. Shaded area, standard deviation across biological replicates. (B) Polar localization index and (C) asymmetry index measurements of ChpA- and phosphorylation-dependent localization of mNG-PilG. Both deletion of *chpA* and inhibition of phosphorylation through ChpA reduce the polar localization index of PilG as well as polarization. (D) Polar localization profiles of mNG-PilG in Δ*fimL* and Δ*fimL* Δ*chpA* mutant after 2h on surfaces. (E) Polar localization index shows that double deletion of *fimL* and *chpA* completely abolishes polar localization of PilG. Single deletion of *fimL* reduces but does not abolish PilG polar localization and polarization (F). Circles, median of each biological replicate. Vertical bars, mean across biological replicates. For the corresponding mean cell fluorescence, see Supplementary Figure 2B.

To rigorously demonstrate that phosphorylation promotes PilG polarization, we interfered with the ability of ChpA to signal to response regulators. Substituting three residues in the histidine kinase domain (D2086A, D2087A, G2088A) blocks autophosphorylation, resulting in a loss-of-function (LOF) mutant ChpA_LOF_ that is unable to transfer phosphate to PilG and PilH (10,18). While ChpA_LOF_ is unable to transfer phosphate, it still localizes to the poles (Supplementary Figure 3AB). Like Δ*chpA*, ChpA_LOF_ neither twitches nor increases intracellular cAMP levels on surfaces (Supplementary Figure 4B) (10). PilG polar localization and asymmetry indices are much lower in ChpA_LOF_ compared to WT (Figure 2BC), showing that ChpA promotes PilG polarization by phosphorylation. ChpA_LOF_ may also weakly bind PilG independent of phosphorylation as PilG polar localization is slightly higher in ChpA_LOF_ compared to Δ*chpA* (Figure 2B). We verified whether *chpA* mutations could affect PilG localization due to cAMP-dependent transcriptional changes. We quantified the localization and polarization of mNG-PilG at low and high cAMP levels by respective deletion of the adenylate cyclase gene *cyaB* and phosphodiesterase gene *cpdA*. PilG localization was largely insensitive of cAMP levels (Supplementary Figure 5). Our results therefore support a model in which T4P mechanosensing events at the leading pole stimulate ChpA to phosphorylate PilG to induce polarization.

PilG localization decreases but is not abolished in Δ*chpA* mutants, suggesting there exists an additional polar binding partner. We hypothesized that FimL, a protein that interacts with the polar hub FimV and PilG, could fulfil that role (20). To characterize the impact of FimL on PilG polar localization, we imaged mNG-PilG in a *fimL* deletion mutant. Polar localization and asymmetry of PilG decreased in *ΔfimL* compared to WT (Figure 2EF). PilG polar localization was entirely abolished in a *ΔfimL ΔchpA* double deletion background (Figure 2E) showing that FimL and ChpA simultaneously recruit PilG to the poles.

Taken together, our results specify the molecular mechanisms driving PilG polar localization and polarization. FimL maintains at least non-phosphorylated PilG at the poles (20). ChpA recruits non-phosphorylated PilG to transfer phosphate, thereby polarizing PilG and drive forward twitching (9).

### Polar recruitment of PilH depends on ChpA but not on its kinase activity

By analogy with PilG, phosphorylation by ChpA could explain the slow increase in PilH polar localization during surface association (Figure 1D). To test this possibility, we quantified mNG-PilH localization in Δ*chpA* and ChpA_LOF_. We first controlled potential indirect effects of cAMP. In a Δ*cyaB* background with low cAMP, PilH polar localization was reduced compared to WT (Supplementary Figure 6C). Conversely, polar localization was stronger in Δ*cpdA* with high cAMP levels. The difference in polar localization between WT and Δ*cpdA* levelled out after 2h of surface contact, as both polar localization indices became indistinguishable (Supplementary Figure 6C). PilH localization is therefore sensitive to cAMP levels. To characterize the additional role of ChpA in PilH localization independently of cAMP, we performed all subsequent mNG-PilH localization experiments in a Δ*cpdA* mutant background (Figure 3).

**Figure 3:**
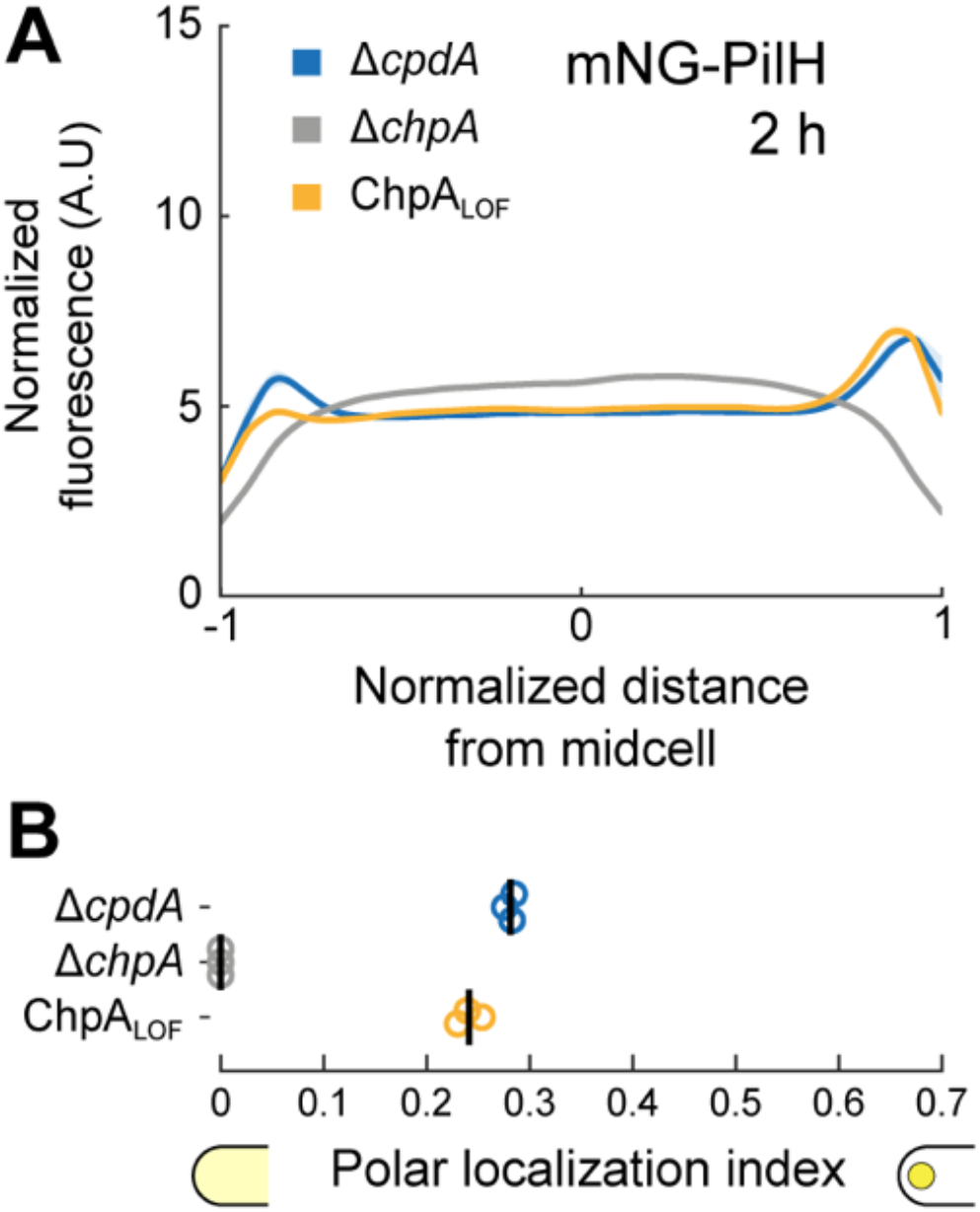
ChpA promotes PilH localization in a phosphorylation-independent manner. (A) mNG-PilH fluorescence profiles in *chpA* mutants after 2h surface growth. To avoid negative effects of low cAMP level on localization of PilH, *cpdA* was deleted in all displayed strains to rescue cAMP level to WT levels (cf. Supplementary Figure 4B). Solid lines, mean normalized fluorescence profiles across biological replicates. Shaded area, standard deviation across biological replicates. (B) PilH polar localization is abolished in Δ*chpA*. PilH polar localization is however maintained in ChpA_LOF_. Circles, median of each biological replicate. Vertical bars, mean across biological replicates. For the corresponding asymmetry index and mean cell fluorescence see Supplementary Figure 2C.

mNG-PilH polar localization is abolished in Δ*chpA ΔcpdA* (Figure 3B), despite nearly identical protein levels as in Δ*cpdA* (Supplementary Figure 2C) and high cAMP levels (Supplementary Figure 4B). Polar localization of PilH is therefore absolutely ChpA-dependent. To address whether PilH localization depends on phosphorylation, we imaged mNG-PilH in the non-functional ChpA_LOF_ mutant with rescued cAMP level *(ΔcpdA* background). The polar localization index of ChpA_LOF_ Δ*cpdA* is close to the one of Δ*cpdA* (Figure 3B). These results show that despite depending on ChpA, polar recruitment of PilH does not depend on phosphorylation through ChpA. It is therefore possible that both non-phosphorylated and phosphorylated forms of PilH localize to the poles. We found that cAMP levels partially control polar localization of PilH. The slow increase in cAMP levels during surface contact (8) could thereby explain the increasing polar localization of PilH over time (Figure 1D).

### The signaling hierarchy between PilG and PilH impacts twitching

The dynamic polarization of PilG during motility and localization of PilH during surface adaptation motivated us to investigate the role of phosphorylation during twitching. In response to mechanosensing, Chp controls forward twitching by polarizing PilB while still enabling spontaneous and collision-induced reversals (9). PilG is necessary for forward twitching while PilH is for reversals. We tracked hundreds of isolated motile cells twitching at the interface between agarose and a glass coverslip (Figure 4AB). We computed their linear displacements which we display as spatial-temporal diagrams (Figure 4C). We had previously shown that *pilG* deletion mutants constitutively reverse, while *pilH* deletion mutants always move forward and never reverse (Figure 4C i and ii) (9). Here, we found that Δ*chpA* and ChpA_LOF_ mutants constantly reversed twitching direction (Figure 4C iii and iv), recapitulating the hyper-reversing phenotype of Δ*pilG*. The inability to phosphorylate PilG and PilH thus phenocopies Δ*pilG*, not Δ*pilH*. Therefore, phosphorylation of at least PilG is required to induce persistent forward twitching.

**Figure 4:**
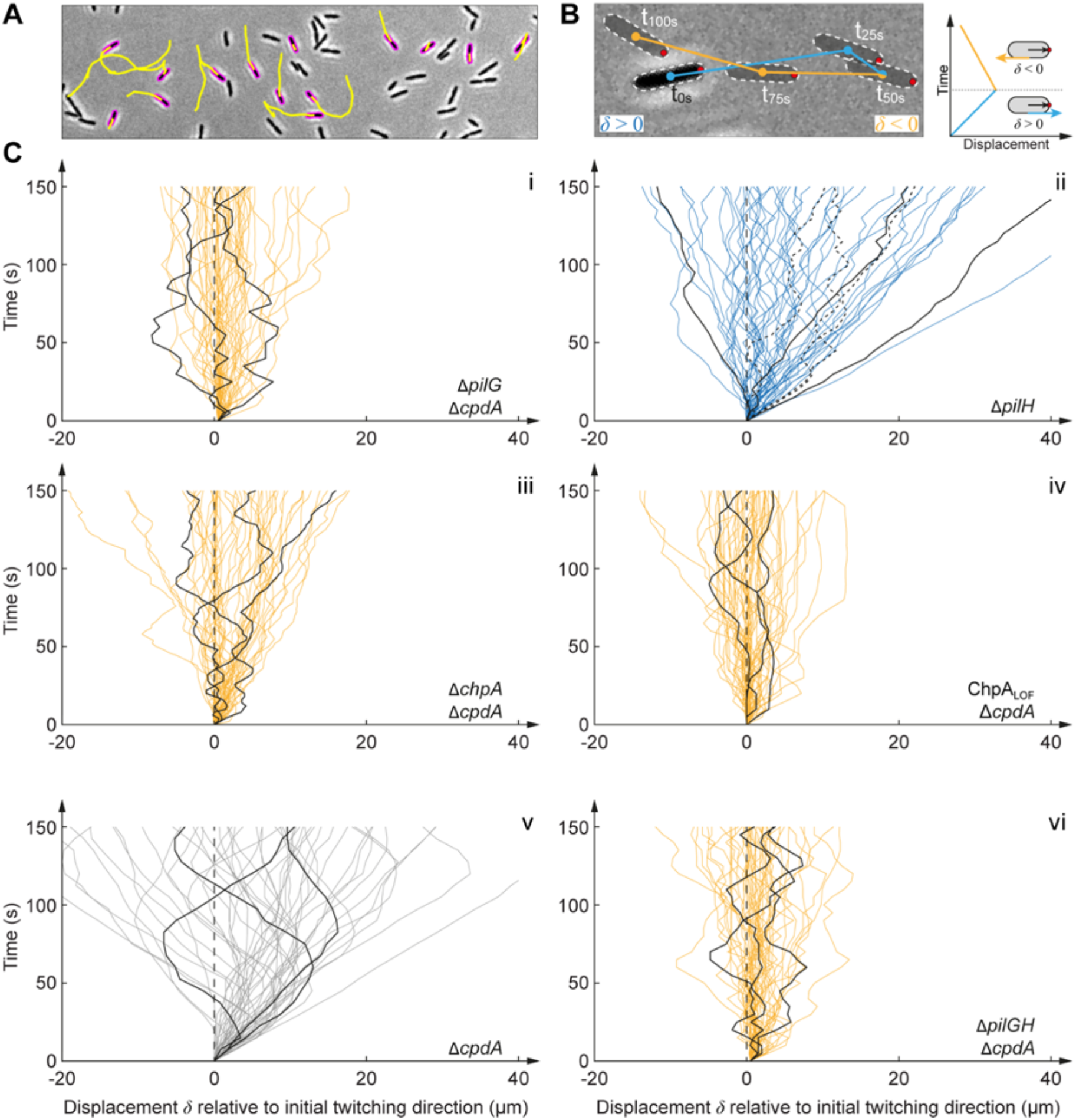
ChpA regulates forward and reverse twitching. (A) Snapshot of 5 min-long twitching trajectories (yellow lines). (B) Computation of displacement maps. A change in the orientation of the track relative to the initially leading pole (red dot) corresponds to a reversal. (C) Displacement maps derived from 50 randomly-selected cell trajectories. Each curve corresponds to an individual trajectory. In each graph, we highlighted three representative tracks for clarity. All graphs correspond to a strain background with high cAMP levels, if necessary by deleting *cpdA*. We use Δ*cpdA* as reference. Deletion and loss-of-function mutation of *chpA* show a hyper-reversing phenotype, similar to deletion of *pilG*. Deletion of *pilH* leads to the opposite phenotype, straight twitching without reversals. Δ*pilG* Δ*pilH* double mutants also hyper-reverse, phenocopying Δ*pilG* and Δ*chpA*/ChpA_LOF_ mutants.

We wondered whether PilH itself was required to trigger reversals in mutants that are already hyper-reversing. The double deletion mutant Δ*pilG ΔpilH* also hyper-reverses, phenocopying Δ*pilG* (Figure 4C vi). This demonstrates that PilH is not inherently required to reverse. PilH may only be required for reversals when PilG is activated by phosphorylation, *i.e*. when cells are already polarized. Our results suggest that PilH does not trigger reversals by directly inhibiting the recruitment of the extension motor PilB independently of PilG (cf. (9)). PilH may rather interfere with the positive feedback from PilG on PilB recruitment and thus on T4P extension. This suggests PilG is the main output regulator of the Chp phosphorylation cascade, controlling directionality of twitching, and that PilH functions antagonistically by mitigating the positive effect of PilG on single-cell mechanotaxis.

### PilG does not directly regulate PilH localization upon surface contact

Twitching motility results suggest that PilH mitigates mechanically-induced PilG activation. To further refine the mechanisms of their antagonistic relationship, we investigated how they impact each other’s localization. We could not distinguish a change in mNG-PilH localization in a Δ*pilG* mutant background, supporting that PilG does not regulate PilH activation (Figure 5AB). Since PilH localization is sensitive to cAMP levels, we confirmed PilG-independent localization in Δ*pilG ΔcpdA* whose cAMP levels are rescued to near WT levels (Figure 5CD).

**Figure 5:**
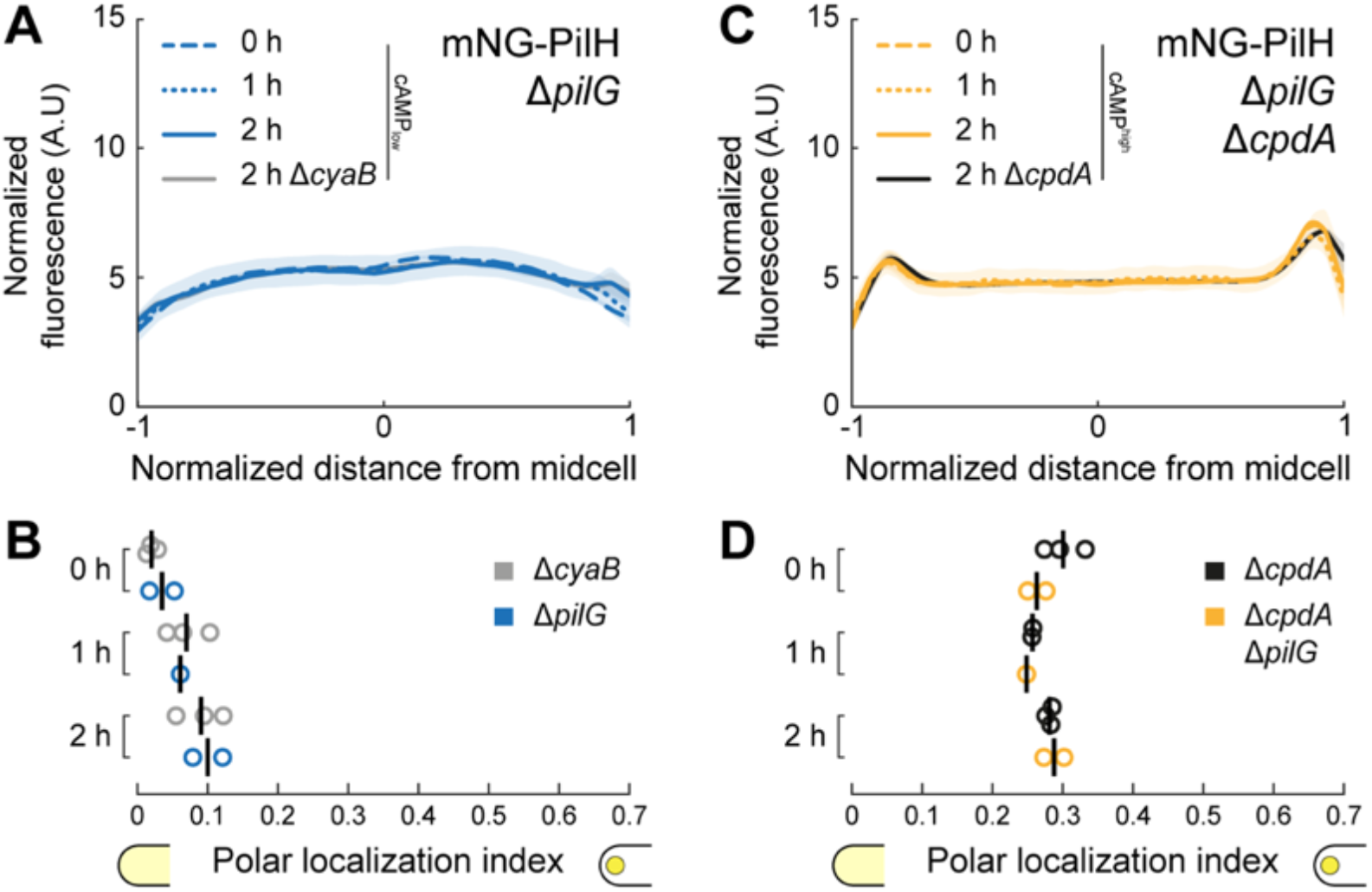
Localization of PilH is not directly affected by PilG. (A) mNG-PilH fluorescence profiles in Δ*pilG* after 2h surface growth (low cAMP). (B) Corresponding polar localization index of mNG-PilH in Δ*pilG*. Recruitment of PilH to the poles is slow but detectable in Δ*pilG*, almost identical to Δ*cyaB* which has similar cAMP levels (Supplementary Figure 4A). (C) mNG-PilH fluorescence profiles in Δ*pilG ΔcpdA* after 2h surface growth (rescued cAMP). (D) Corresponding polar localization index of mNG-PilH in Δ*pilG ΔcpdA*. PilH polar recruitment is indistinguishable from Δ*cpdA*, which also has elevated cAMP levels (Supplementary Figure 4A). Solid lines, mean normalized fluorescence profiles across biological replicates. Shaded area, standard deviation across biological replicates. Circles, median of each biological replicate. Vertical bars, mean across biological replicates. For the corresponding asymmetry index and mean cell fluorescence see Supplementary Figure 2D.

### PilH activation controls PilG polarization

PilH may function antagonistically by directly inhibiting PilG. To characterize how PilH impacts PilG polarization, we quantified PilG localization in Δ*pilH*. Polar localization of PilG is stronger in Δ*pilH* relative to WT and to Δ*cpdA* whose cAMP levels nearly match Δ*pilH* (Figure 6AB) (9). In addition, the asymmetry index of mNG-PilG is larger in Δ*pilH* than in WT (Figure 6C). We conclude that PilH represses PilG polar localization and polarization.

**Figure 6:**
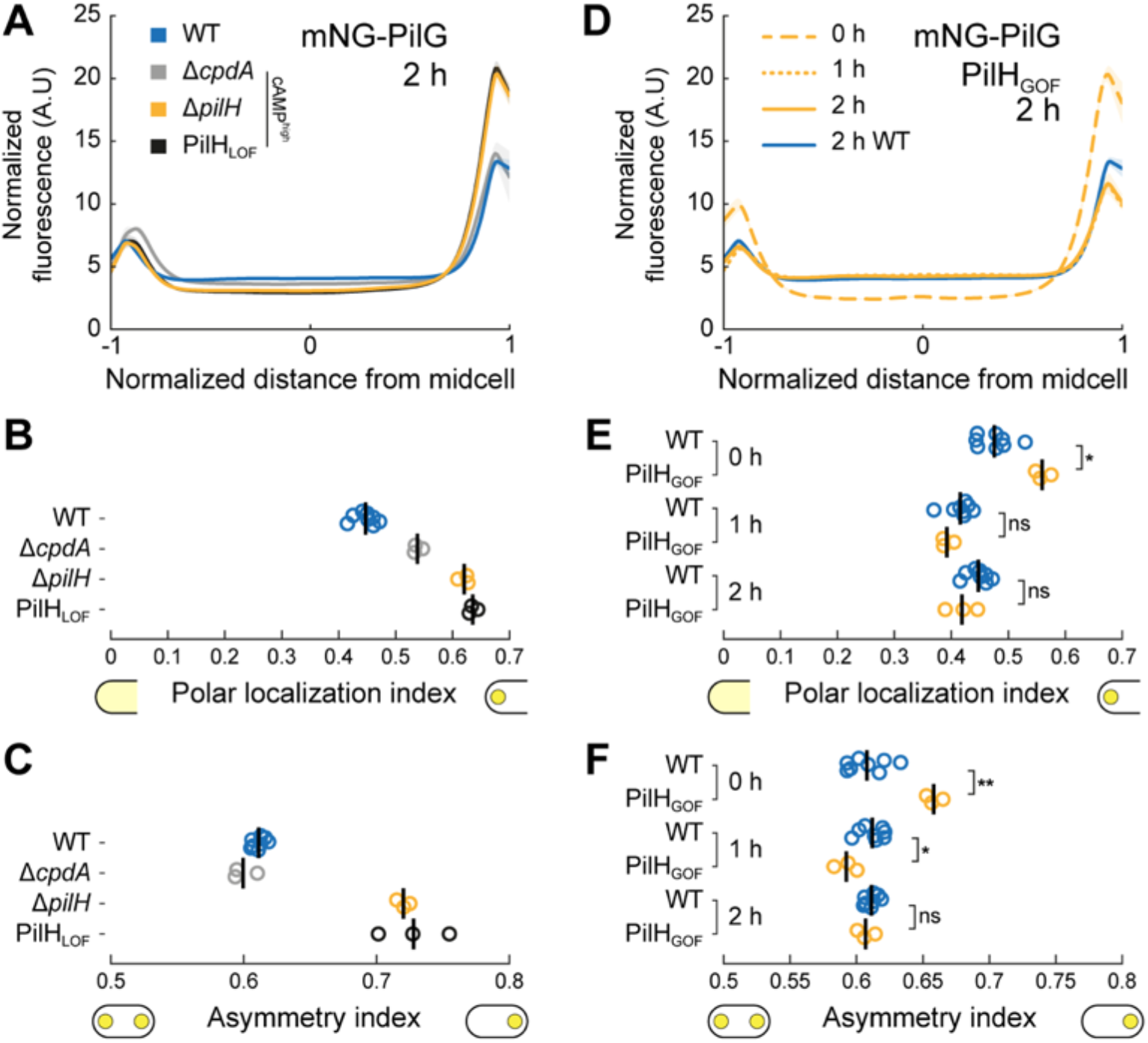
PilH activation modulates PilG polar localization. (A) mNG-PilG fluorescence profiles in *pilH* mutants after 2h surface growth. We included Δ*cpdA* for comparison since both *pilH* mutations result in high cAMP levels. PilG is more polar and asymmetric in Δ*pilH* and PilH_LOF_. (B) Polar localization index and (C) asymmetry index measurements of PilH-dependent localization of mNG-PilG. (D) Time course of PilG localization in PilH_GOF_ on surface. For clarity only the 2h profile of PilG in wild type was included. In PilH_GOF_, polar localization of PilG is comparable to Δ*pilH* and PilH_LOF_ at 0h, but similar to wild type after 1h and 2h surface growth. (A and D) Solid lines, mean normalized fluorescence profiles across biological replicates. Shaded area, standard deviation across biological replicates. (B, C, E, F) Circles, median of each biological replicate. Vertical bars, mean across biological replicates. *, p<0.05; **, p≤0.001; ns, not significant. For the corresponding mean cell fluorescence see Supplementary Figure 2E. For corresponding example micrographs see Supplementary Figure 1B.

We wondered whether PilH activation by ChpA is necessary to repress PilG polarization. To characterize the effect of PilH phosphorylation on PilG localization, we generated a PilH mutant that cannot be phosphorylated by substituting the catalytic aspartate 52 with alanine to make PilH_LOF_ (LOF, loss-of-function). PilH_LOF_ cells twitch forward without reversing and have elevated cAMP levels similar to Δ*pilH* mutants, confirming the loss-of-function mutation (Supplementary Figure 7). In PilH_LOF_, PilG-mNG polar localization index and asymmetry index are indistinguishable from Δ*pilH* (Figure 6BC). Substituting aspartate 52 to glutamate yields a gain-of-function (GOF) mutant, PilH_GOF_, which is predicted to adopt an active phosphorylated conformation but loses the ability to be phosphorylated. cAMP levels and reversal rate measurements confirmed that PilH_GOF_ is functional and can complement a deletion of *pilH*, although both phenotypes are not quantitatively restored to WT levels (Supplementary Figure 7). In PilH_GOF_, PilG is more polar and more asymmetric than WT at 0h (Figure 6D-F). Both indices however decrease over time on surface to eventually reach WT levels. This is consistent with a model where PilH in its active conformation reduces the polar localization of PilG and thus mitigates PilG polarization. This effect is independent of PilH-related phosphate flow since PilH_GOF_ cannot get phosphorylated. In addition, this only takes effect after surface contact and is not an inherent property of PilH_GOF_.

### ChpA recruits PilH_GOF_ to the poles upon independently of phosphorylation

Since ChpA cannot phosphorylate PilH_GOF_, we wondered whether polar recruitment of PilH_GOF_ played a role in mitigating the local positive feedback instilled by PilG. Like wild-type PilH, PilH_GOF_ slowly increases polar localization (Figure 7AB). PilH_LOF_ also slowly localizes to the poles but less pronounced than PilH or PilH_GOF_ (Supplementary Figure 9). The slow recruitment in all three mutants is consistent with a scenario where polar localization of PilH is independent of phosphorylation. Like for wild-type PilH, polar localization of PilH_GOF_ is ChpA-dependent but independent of ChpA’s ability to autophosphorylate and transfer phosphate (Figure 7CD). This suggests that ChpA recruits both PilH and activated PilH-P to the poles. A direct interaction between PilH forms and ChpA has yet to be clearly demonstrated.

**Figure 7:**
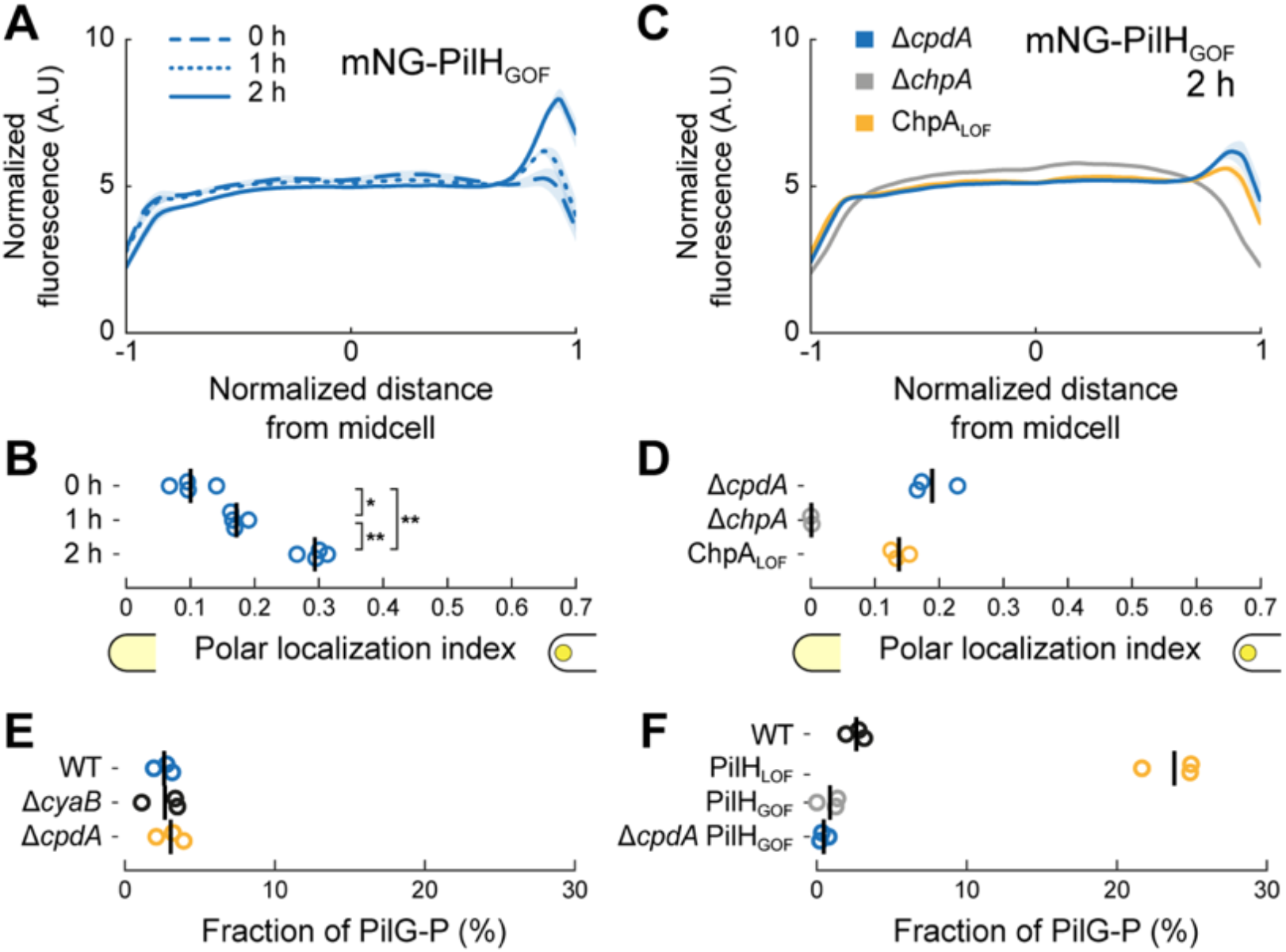
ChpA recruits active PilH to the pole in a phosphorylation-independent manner. (A) Time course fluorescence profiles of mNG-PilH_GOF_. (B) Like wild-type PilH, PilH_GOF_ gets recruited to the poles over time on surface. C) Fluorescence profiles of mNG-PilH_GOF_ after 2h on surface in *chpA* mutants. (D) Polar localization index measurements show polar localization of mNG-PilH_GOF_ depends on ChpA, but not on its kinase activity. (A, C) Solid lines, mean normalized fluorescence profiles across biological replicates. Shaded area, standard deviation across biological replicates. (B, D) Circles, median of each biological replicate. Vertical bars, mean across biological replicates. *, p<0.05; **, p≤0.001; ns, not significant. For the corresponding asymmetry index and mean cell fluorescence see Supplementary Figure 2FG. (E) PhosTag™ measurements of the fraction of phosphorylated Flag-PilG (PilG-P) after 2h surface growth in low (Δ*cyaB*) and high (Δ*cpdA*) cAMP levels. (F) Fraction of phosphorylated PilG after 2h surface growth in PilH_GOF_ and PilH_LOF_. To rule out a negative effect of lower cAMP level in PilH_GOF_, we deleted *cpdA* in that mutant. Circles correspond to biological replicates, black bars represent their mean. See Supplementary Figure 8 for a representative PhosTag™ gel.

In summary, PilH is required for proper localization of PilG. Without PilH, PilG localizes exceedingly asymmetrically and likely irreversibly locked to one pole, explaining the unidirectional twitching phenotype observed in Δ*pilH* (9). Upon recruitment to the poles, PilH in its active conformation reduces polar localization of PilG. Since PilH and PilH_GOF_ polar localization are ChpA-dependent, we conclude that PilH functions through ChpA. PilH locked in its active conformation, PilH_GOF_, is sufficient to reduce PilG localization. This process does not require phosphate transfer involving PilH, disproving the hypothesis that PilH acts as a phosphate sink for PilG and ChpA (11,18).

### PilH activation regulates PilG phosphorylation

Since phosphorylation stimulates PilG polarization, PilH could indirectly regulate PilG polarization by controlling PilG phosphorylation. We thus measured the fraction of phosphorylated PilG in whole-cell lysates by PhosTag™ assays in PilH_LOF_ which causes strong polarization, and in PilH_GOF_ which has WT-like polar localization. First, we verified that phosphorylated Flag-tagged PilG migrated as two bands on PhosTag™ gels (Supplementary Figure 8A). The slower migrating band was absent in PilG point mutants that cannot be phosphorylated (20). Since PilH_LOF_ and PilH_GOF_ mutants affect cAMP production we verified that cAMP levels do not affect phosphorylation of PilG by measuring PilG-P in WT, Δ*cyaB* (low cAMP) and Δ*cpdA* (high cAMP) (Figure 7E). In PilH_LOF_, the fraction of phosphorylated PilG is higher than in WT (Figure 7F). The fraction of phosphorylated PilG-P is relatively reduced in PilH_GOF_ and in PilH_GOF_ Δ*cpdA* (Figure 7F). This suggests PilH in its phosphorylated conformation reduces PilG phosphorylation without the need of phosphate flow to or from PilH.

## Discussion

How cells establish and switch polarity are critical questions in biology. Polarity is an essential requirement of the physiology of many bacteria. For example, cells polarize by asymmetric localization of cellular components during motility or asymmetric division *(21)*. Our results bring a high-resolution perspective on the Pil-Chp signaling network controlling reversible polarity in *P. aeruginosa* in response to mechanosensing (Figure 8). In particular, we identified the role of PilG and PilH activation in relaying information from mechanical input to a motility response. Cells sense surface contact through T4P. During attachment and retraction, T4P activate Chp through the receptor PilJ *(7,8)*. PilJ stimulates ChpA autophosphorylation at that pole. ChpA then recruits PilG to transfer phosphate, thereby polarizing the cell. This in turn stimulates T4P extension via the recruitment of at least FimX and PilB *(9)*. Increased T4P extension locally stimulates mechanosensing at that same pole, feeding a local positive feedback, which stabilizes polarization and forward motility. We showed that PilG phosphorylation by ChpA induces polarization. This supports a model in which local activation by T4P triggers and stabilizes polarization, establishing the most direct molecular link between mechanical input (T4P retraction on surface) and cellular output (PilG phosphorylation). Setting polarity only requires PilG, supporting the hypothesis that PilG is the main output regulator of the Chp system *(10,11,18)*.

**Figure 8:**
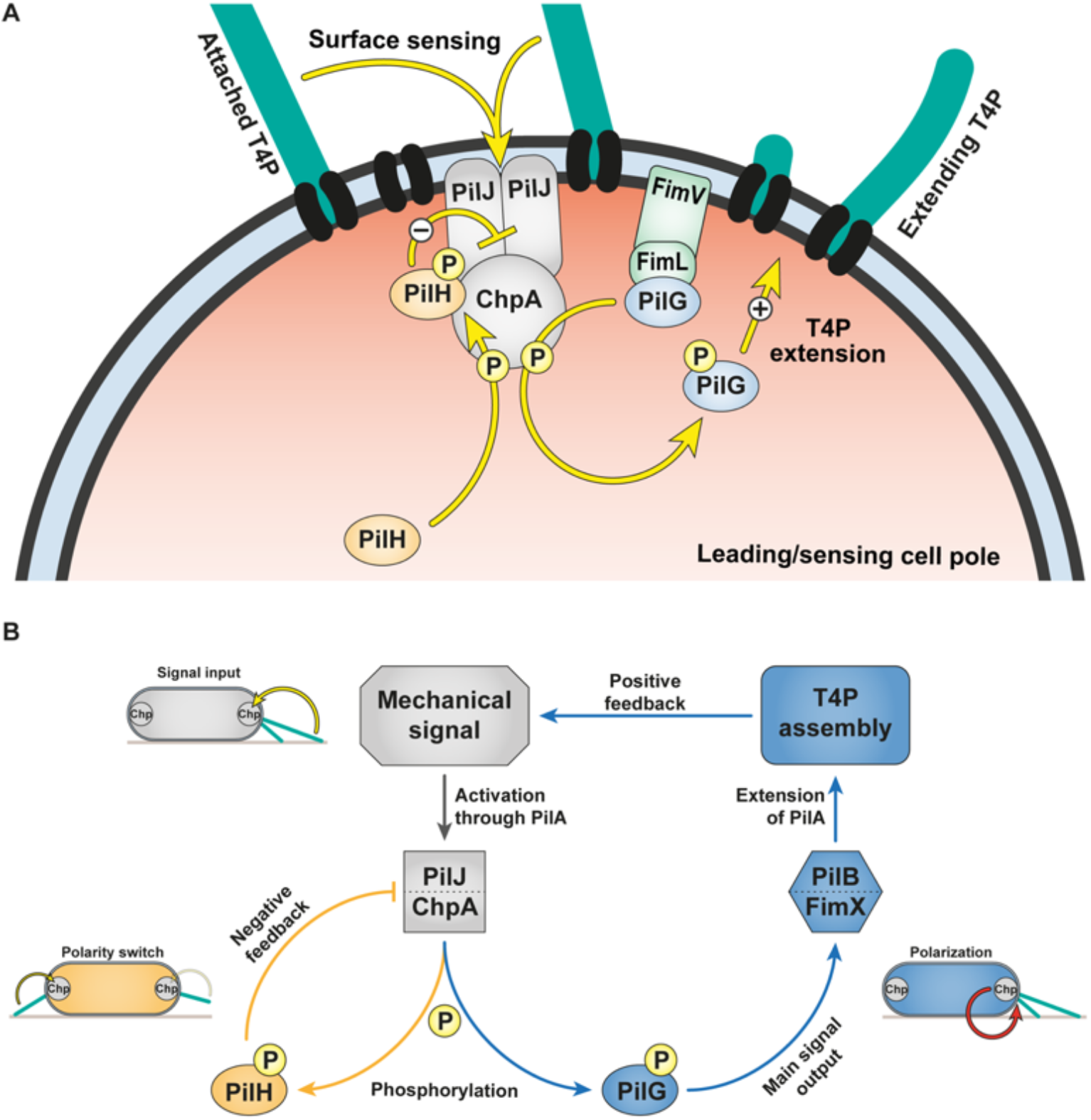
A model of signalling feedback in *Pseudomonas aeruginosa* mechanotaxis. (A) Model of phosphate flow through Chp at the leading pole of twitching cells. PilG is recruited by two polar proteins, ChpA and FimL. FimL maintains PilG close to Chp to restrict diffusion. ChpA phosphorylates PilG so induce polarization, which recruits FimX and PilB to promote T4P extension (9). Polar localization of PilH is independent of phosphorylation. However, a conformational change induced by phosphorylation is required to activate PilH. PilH-P mitigates the polarizing effect of PilG by reducing phosphorylation. (B) Overview of the Pil-Chp regulation circuit. Chp senses a mechanical signal when T4P attach at the leading pole of motile cells. Surface sensing involves interaction of T4P filament monomers PilA and Chp’s receptor PilJ (7,8). As a consequence, ChpA becomes more active and phosphorylates PilG. PilG constitutes the main signal output of Chp, eventually polarizing cells by recruiting FimX and PilB to the actively sensing pole where they locally activate T4P extension. As T4P themselves are the surface sensor, this creates a positive feedback loop. ChpA also phosphorylates PilH to balance the main signal output by modulating PilG’s activity and localization. PilH achieves this by decreasing PilG phosphorylation, potentially by inhibiting PilJ/ChpA. Arrows depict hypothesized functional interactions, not direct protein-protein interactions.

Local activation of PilG alone cannot explain asymmetric activation of T4P extension. In *E. coli*, CheY-P quickly diffuses to flagellar basal bodies located throughout the cell envelope after phosphorylation (22–24). Conceivably, cells need to restrict diffusion of PilG once phosphorylated to limit the positive feedback to the leading input pole. We postulate FimL, which is maintained at the poles by interaction with the polar landmark protein FimV (20,25), is involved in focusing PilG to the mechanosensing input pole (the leading pole in motile cells). This may restrict PilG-P diffusion towards the opposite pole. Interestingly, the N-termini of FimL and ChpA are homologous, suggesting a connection between the two proteins, for example in interacting with PilG (20). FimL is further involved in phosphorylation-independent asymmetry of PilG, although it is not as pronounced as phosphorylation-dependent asymmetry (Figure 2). While the feedback-driven asymmetry established by phosphorylation of PilG enables dynamic switching of polarity in response to mechanical signal input, the role of FimL-dependent localization of PilG remains to be clarified in future studies.

Local positive feedback promotes forward locomotion but prevents directional changes. Without means to counteract this feedback, motile cells would trap themselves in corners, and cell groups would jam during collective twitching (9,26). *P. aeruginosa* avoids locking itself into one polarization by mitigating the positive feedback. At the molecular level, the response regulator PilH counteracts PilG to invert polarization and enable reversals of twitching direction. Multiple conflicting molecular models have been proposed for the antagonizing effect of PilH (10,11,18). We here resolved the mechanism of PilH function in the Chp signaling system. We show PilH directly regulates PilG polarization by reducing PilG phosphorylation. To reverse PilG polarization, PilH must transition to its active conformation, for which it most likely must be phosphorylated. PilH-P may diffuse across the cytoplasmic space to create a global antagonizing effect at both poles. This in turn interrupts the positive feedback sustained by PilG at the leading pole. PilH and PilH-P localize to the poles without polarizing, making the investigation of PilH dynamic activation in motile cells complex. Both PilH and constitutively-activated PilH_GOF_ require ChpA for polar localization. This indicates that PilH interacts with ChpA, and that once phosphorylated PilH-P inhibits PilG phosphorylation by ChpA, potentially by blocking or inhibiting the phosphotransfer domain(s) responsible for PilG phosphorylation. PilH-P may also stimulate the receiver domain of ChpA (ChpA_REC_) that may function as a phosphate sink for PilG. The autodephosphorylation rate of ChpA_REC_-P is order of magnitudes faster than of PilG-P or PilH-P, consistent with a sink function (18).

Polar localization of PilH gradually increases during surface adaptation, however, independently from PilH phosphorylation. Instead, polar recruitment of PilH correlates well with cAMP levels (cf. Supplementary Figure 4A and Supplementary Figure 6C), suggesting PilH localization is regulated by transcription (6,27). However, PilH localization doesn’t correlate with mNG-PilH protein abundance (Supplementary Figure 6C vs E). It is therefore unlikely that protein expression alone explains PilH polar recruitment. As both PilH and PilH_GOF_ require ChpA for polar localization, the factors responsible for recruiting PilH to the pole may at least involve interaction with ChpA. By contrast, PilG polar localization and polarization are not considerably affected by surface sensing on long timescale, indicating some level of homeostasis. We also found that PilG is unexpectedly polarized in cells taken right out of liquid culture conditions, but this might be due to limitations of visualization (Figure 1D).

Chp-like systems with multiple response regulators may be involved in surface-induced and taxis-related phenotypes in other species. Chp homologs are present in a number of bacterial species, for example many gamma-proteobacteria including *P. aeruginosa, Acinetobacter baumanni, Stenotrophomonas maltophilia* or *Dichelobacter nodosus* among others (20). In *Acinetobacter* species T4P mediate surface motility and natural transformation. ChpA and PilG are required for motility and transformability in *A. baumanii* (28) and motility in *A. baylyi* (29). PilH may also function antagonistically to PilG as in *P. aeruginosa* (28).

The cyanobacterium *Synechocystis* uses phototaxis to direct T4P-dependent surface motility. Several chemotaxis-like systems are involved, three of which comprise two response regulators as in Chp (30). The response regulators PixG, PilG and TaxP2 have an N-terminal PATAN domain that is missing in PilG of *P. aeruginosa* and related systems. The PATAN domain mediates direct interaction with the T4P extension motor PilB whose localization sets twitching direction in response to light (30–32). The output functions of their second response regulators PixH, PilH and TaxY2 have yet to be resolved, but based on our results in *P. aeruginosa*, one could anticipate that they counter the function of their cognate first response regulator. In *Myxococcus xanthus* a chemotaxis-like system called Frz regulates the frequency of motility reversals, in the same manner as Chp in *P. aeruginosa*. Frz controls the localization of MglA, MglB and RomR to the leading or lagging pole, thereby setting the direction of gliding and twitching motility (33–37). Here, the output response regulators FrzX and FrzZ show clear localization to opposite poles, in contrast to PilG and PilH (35).

Although Chp comprises a more complex signaling network than the dominant model of chemotaxis systems, the Che system controlling flagellar rotation in *E. coli*, there are even more complex chemosensory systems in many species. They often comprise several copies of chemosensory components, especially response regulators, but also multiple sets of full chemotaxis systems. Why some species require such complexity in chemotaxis systems remains poorly understood. In *Rhodobacter sphaeroides* multiple chemotaxis systems control flagellar rotation in response to different signal inputs (38). Remarkably, only one main response regulator actively stops the flagellar motor, while others may only modulate the effect of the main regulator, indicating a shared signaling principle with Chp (39).

We here decoded the signaling pathway of the Chp system during mechanotaxis. Our results resolve the molecular mechanisms of signaling with two response regulators. At the same time, our model proves a rationale for the complexity of Chp by demonstrating the necessity of each component in setting a local positive feedback while still enabling reversals. Our data therefore provides a new framework to understand more complex sensory systems in bacteria.

## Materials and Methods

### Bacterial Strains, growth conditions and media

For all experiments *P. aeruginosa* PAO1 ATCC 15692 (American Type Culture Collection) was used. For vector constructions and conjugative mating *E. coli* strains DH5α and S17.1 were used, respectively. All bacterial strains were grown in LB medium (Carl Roth) at 37 °C with 280 rpm shaking. Regular solid LB media were prepared by adding 1.5% (wt · vol^-1^) agar (Fisher Bioreagents) and appropriate antibiotics, for selection of *E. coli* 100 μg.ml^-1^ ampicillin or 10 μg.ml^-1^ gentamycin, for selection of *P. aeruginosa* 300 μg.ml^-1^ carbenicillin or 60 μg.ml^-1^ gentamycin. For twitching and protein localization experiments semi-solid tryptone media were prepared by autoclaving (5 g.l^-1^ tryptone (Carl Roth), 2.5 g.l^-1^ NaCl (Fisher Bioreagents), 0.5 % (wt.vol^-1^) agarose standard (Carl Roth)). For measurements of cAMP levels on solid surfaces LB plates containing 1% standard agarose were prepared by autoclaving (for the PaQa reporter), or regular 1.5% LB agar plates were used (for the PlacP1 reporter). Surface growth for PhosTag™ assays was carried out on regular 1.5% LB agar plates.

### Strains and vector construction

Strains are listed in Supplementary Table 1, plasmids and corresponding oligonucleotides are listed in Supplementary Tables 2 and 3, respectively. *Pseudomonas aeruginosa* mutants were generated as described previously (9). Genes were deleted or integrated by two-step allelic exchange according to (40) using the suicide vectors pEX18_Amp_ or pEX18_Gent_. For genomic in-frame gene deletions, approximately 500 to 1000 base pair fragments of the up-and downstream regions of the designated gene were combined by PCR amplification and subsequent Gibson assembly (41). Marker-free deletions were verified by PCR and sequencing. In-frame insertions were generated essentially the same way. Substitutions of wild-type genes with mutated genes (e.g. point mutants) were integrated into the corresponding deletion strains or wild type. Proteins were fluorescently labelled typically by N-terminal fusion separated by a 5xG linker, and integrated into the wild-type gene locus. Functionality was tested by monitoring single cell twitching and measurements of cAMP levels (see also (9)). Plasmids were constructed using standard Gibson assembly protocols (41) and introduced into *P. aeruginosa* cells by conjugative mating with *E. coli* S17.1 as donor.

### Fluorescence microscopy

Microscopy was performed on an inverted Nikon TiE epifluorescence microscope using NISElements (version AR 5.02.03). For phase contrast microscopy a 40× Plan APO NA 0.9 phase contrast objective was used. For fluorescence microscopy a 100× Plan APO NA 1.45 phase contrast oil objective and Semrock YFP-2427B or TxRed-A-Basic-NTE filters were used as needed. Microscope settings were kept consistent throughout all experiments to ensure comparability of fluorescent intensities, unless stated otherwise. Fluorescent images were background subtracted and snapshots and movies were generated with ImageJ (version 1.53). Data were analyzed with custom scripts using Python (version 3.8.5) and MATLAB (version R2019b), as specified in detail below. Custom codes are available on Github (https://github.com/PersatLab/Mechanotaxis).

### Media and cell preparation for single cell twitching experiments

Plates were prepared by autoclaving tryptone medium supplemented with 0.5% agarose, cooling to 70 °C in the autoclave followed by cooling to 55 °C for 20 min in a water bath. 28 ml medium was poured into 90 mm petri dishes and dried in a flow hood for 30 min. Plates were always stored for one day at 4 °C in a plastic bag and used the next day or maximum after two days. Exponentially growing cells (filtered LB medium, OD_600_ = 0.2-0.8) were diluted to OD_600_ = 0.2. After prewarming the plates for 45-60 min a 16 mm round pad was cut out and 1 μl of the diluted cell suspension was pipetted onto the upper side of the agarose pad (i.e. the side that was not in contact with the plastic dish bottom). The pads were immediately flipped onto a microscope glass bottom dish (P35G-1.5-20-C, MatTek) and 4 droplets of PBS were added at the sides without touching the pad to prevent drying. The dishes were used for imaging immediately or incubated at 37 °C for later imaging.

### Quantification of protein localization

Cells were prepared and imaged as described above. Although protein localization was analyzed independent of twitching direction, motile cells were imaged. Motile cells are typically visible after 1 and 2h but not at 0h. The analysis was carried out essentially as described previously (9). Fluorescent images of mNeonGreen fusion proteins were acquired at 0, 1 and 2h after preparation of the dishes. 0h samples were taken within 1-5 minutes after the cell suspension was pipetted onto the tryptone agarose pads. Imaging settings (objective and filters, excitation power and time, binning) were kept constant across all experiments to ensure comparability of the fluorescent signal. Cells were segmented using phase contrast images and fluorescence profiles were extracted with BacStalk (version 1.8, (42)). Typically, several dozens to hundreds of cells were segmented per replicate. Fluorescent profiles correspond to the mean pixel value of a transversal section of the cell along the mid-cell axis (see also (9)). For comparison of proteins with different expression levels the profiles were normalized by the total fluorescence of the cell and rescaled the cell length. Cells were oriented so that the dim pole is at x = −1, the bright pole at x = 1, and mid-cell at x = 0. Mean profiles and standard deviations were computed for every biological replicate. We defined a circular polar area according to the ratio between cell width and cell length to account for differences in cell size. To quantify the extent of polar localization vs cytoplasmic localization we computed a polar localization index. As baseline profile corresponding to a polar localization index of 0 the fluorescence profile of soluble mNeonGreen expressed from pJN105-mNG plasmid (uninduced) was determined. We subtracted the average mNG baseline profile plus standard deviation from the measured profiles. The remaining integrated signal at the defined polar areas divided by the total polar fluorescence corresponds to the polar localization index. Values below zero can occur with this correction for non-polarly localizing proteins due to noise. In this case, the polar localization index was set to 0. The baseline correction was applied because the profiles of exclusively cytoplasmic localizing proteins are not flat due to decreasing signal intensity towards the poles that results from the geometry of the cell poles (see profiles of non-polarly localizing proteins such as mNG-PilH in Δ*chpA*). A polar localization index of 0 corresponds to completely cytoplasmic proteins whereas a value toward 1 corresponds to polarly localized proteins. Due to the applied correction method values of exactly 1 can never be reached, even for proteins that would localize exclusively at the poles. The correction method applied here differs from the method we used previously (9) because we found it to be more accurate for proteins clearly localizing to the poles but to a low extent. Therefore, polar localization indices show here can’t be compared directly to the indices published in (9).

We similarly computed an asymmetry index (previously called symmetry index (9)) by taking the ratio between the maximum polar total fluorescence (comparing the total fluorescence of opposite polar areas) and the sum of the polar total fluorescence (total fluorescence of both polar areas). A value of 0.5 corresponds to a perfectly symmetric bipolar localization whereas a value of 1 corresponds to a perfectly unipolar (asymmetric) localization.

We generated average localization maps of whole cells using the demograph function of BacStalk. We selected 42 cells with similar length around 4 μm. The cell lengths and fluorescent intensities were normalized and subsequently averaged using a custom ImageJ script. Note, these average maps only give a rough visual representation of protein localization and do not include all data that was acquired and used to generate quantitative localization indices as described above.

### Spatiotemporal cumulative displacement maps

Cells were prepared and imaged as described above. The analysis was carried out as described previously (9). Movies of single twitching cells were recorded by phase contrast microscopy at 0.2 frames per second for 5 min at room temperature. Movies were processed with a custom ImageJ macro to ensure compatibility with the downstream analysis. Briefly, movies were cut into quadrants, drift was corrected if necessary using the StackReg plugin (version July 7, 2011; (43)), and image sequences were saved as individual tif images. BacStalk (version 1.8, (42)) was used to segment and track cells. Cell tracks were analyzed with a custom MATLAB script. Briefly, we defined a cell orientation unit vector t^→^ from the center of mass (CM) to the initial leading pole. We determined the initial leading pole by comparing the scalar products of the unit vectors from CM to poles A and B (arbitrary classification) to the normalized displacement vector d^→^ in the first frame in which the cell was classified as moving (speed threshold: one pixel per frame for at least three consecutive frames). We determined the cell displacement δ relative to the initial leading pole using vectors t^→^ and d^→^. δ is positive if cells twitch forward and negative if they reverse and twitch backward. We generated spatiotemporal displacement maps by cumulating the direction-corrected displacement as a function of time.

### Reversal frequency of isolated cells

Cells were prepared and movies recorded as described above. The analysis was carried out as described previously (9). Briefly, for each frame (starting from the first frame in which the cell was classified as moving) the scalar product between the normalized displacement vector d^→^ and the cell orientation unit vector t^→^ was determined and rounded. This yielded a series of numbers that correspond to movement of the cell toward the initial leading pole (1), toward the initial lagging pole (−1) or no movement (0) at that timepoint (timepoints with no movement were removed). Cells were considered moving in the same direction (relative to the initial leading pole) as long as the sign remained the same, and counted reversing when a change of sign occurred. Reversals were only considered if at least two subsequent frames before and after the reversal had the same sign (to correct for frequent sign changes when a cell was close to non-moving). To calculate the reversal frequency we divided the sum of all considered reversals by the total tracked time over all cell tracks for each biological replicate.

### Reversal frequency after collisions

Cells were prepared and movies recorded as described above. The analysis was carried out as described previously (9). We counted cell-cell collisions and potential subsequent reversals manually using the ImageJ plugin Cell Counter. A collision was only considered if the cell was moving for at least three frames in the same direction prior to the collision and the collision lasted for at least two frames (frame interval 5 sec). Collisions with angles below roughly 20° were not considered. We considered a reversal following a collision only if it occurred within five frames after the collision ended. Freshly divided cells were not considered. We calculated the frequency of reversals following a collision by dividing the sum of all considered reversals by the sum of all considered collisions for each biological replicate.

### Fluorescent labelling of pili

Cells were essentially prepared and imaged as described above, with the exception of staining T4P prior to microscopy and increasing the fluorescence excitation power from 20 to 50%. We used a strain with mutated PilA (PilA_A86C_ (44)) expressed from its chromosomal native locus. Prior to preparing the microscope slide, we labelled PilA_A86C_ by supplementing the culture medium with 5 μl of the maleimide-conjugate fluorescent dye Alexa Fluor™ 488 C5 maleimide (Thermo Fischer A10254). Cells were incubated in the dark for 30 min, washed with PBS and used for fluorescence microscopy. Note, only PilA_A86C_ in the extended pili are labelled during the staining process, and more non-labelled proteins get produced after the washing. Therefore, not all pili are visible in the recorded movies, and some cells may still twitch without visible pili.

### cAMP quantification using PaQa-YFP and PlacP1-YFP reporter

Single cell cyclic adenosine monophosphate (cAMP) levels were measured using the PaQa-YFP/PrpoD-mKate2 or PlacP1-YFP/POXB20-mKate2 reporter system as described previously (8). *Pseudomonas* was transformed with reporter plasmid pUCP18-PaQa-YFP/PrpoD-mKate2 and pUC18-PlacP1-YFP/POXB20-mKate2, respectively. PaQa is a native *Pseudomonas* promoter responsive to cAMP, PrpoD is a constitutive promoter. LacP1 is a synthetic promoter responsive to cAMP, OXB20 is a strong constitutive promoter (Oxford Genetics Ltd. (UK), Sigma).

For the PaQa measurements, cells were grown overnight in LB-carbenicillin, diluted to OD_600_ = 0.05 and grown until mid-exponential phase (OD_600_ = 0.4-0.8). After dilution to OD_600_ = 0.1, liquid samples were imaged and in parallel 100 μl of the diluted culture were plated on a 1 % agarose plate (LB, no antibiotics). After 3 h at 37 °C cells were harvested by adding 1.5 ml LB to the plate and gently shaking it. OD_600_ was measured and set to 0.1 for imaging. Images in YFP and mKate2 channels were acquired. Cells were segmented and mean cellular fluorescence was measured using BacStalk (42). We then used a custom Python script to compute median PaQa-YFP to mKate2 fluorescent intensity ratios.

For the lacP1 measurements strains containing the reporter plasmid were grown on LB agar plates with 400 μg/ml Carbenicillin (GoldBio) at 37°C to obtain single colonies. Single colonies were grown in liquid broth in a deep well plate overnight, 37°C, shaking. Overnight cultures were diluted and grown at 37°C for 3 hours. After 3 hours a fraction of the bacteria was fixed by the addition of 4% methanol-free paraformaldehyde (PFA, Thermo Scientific) for 10 min at room temperature and the reaction was quenched by the addition of 0.3 M glycine. A second fraction of bacteria was spotted onto LB/Carbenicilin plates and grown for 2 hours, after 2 hours the spots were scraped into PBS and fixed by the addition of PFA and glycine, as above. Samples were diluted in PBS and analyzed on an LSRII flow cytometer the following day in the UCSF Parnassus Flow CoLab, (RRID:SCR_018206). Data were exported from FlowJo and analyzed using the same custom Python script as above. We reduced the sample size to 1000 cells per replicate by randomizing and reducing the data from each sample separately.

### Phos-tag™ SDS PAGE and immunoblotting to detect phosphorylated PilG in vivo

*Pseudomonas aeruginosa* PAO1 and selected mutant strains expressing chromosomal 3xFlag-PilG were grown on LB agar plates at 37°C to obtain single colonies. Single colonies were grown in liquid LB broth overnight. Overnight cultures were diluted in LB broth and grown at 37°C for 3 hours. After 3 hours a fraction of the bacteria was harvested by centrifugation (10 000g, 5min, 4°C), the pellet was suspended in Laemmli sample buffer (Biorad) with 2.5% 2-mercaptoethanol and 0.1U/ul benzonase (Millipore) and frozen at −20°C. A second fraction was plated on LB plates and incubated at 37°C. After 2 hours bacteria were harvested into 1ml cold buffer (5 mM MgCl_2_ / 50 mM KCl / 50 mM Tris pH 7.5 / 2 mM DTT) and centrifuged at 10 000xg for 5min at 4°C. Supernatants were removed and pellets suspended in cold reducing Laemmli sample buffer (Biorad) containing 0.1U/μl Benzonase (Millipore) and frozen at −20°C. Protein concentration was measured using Pierce™ 660nm Protein Assay Reagent with Ionic Detergent Compatibility Reagent. Two μg of each sample were separated by 12% SDS-PAGE with 100μM Phos-tag™ acrylamide (Wako Chemicals, Richmond, VA, USA) and 200μM MnCl_2_. Phos-tag™ acrylamide gels were electrophoresed at a constant current of 30mA in running buffer (25 mM Tris, 192 mM glycine, 0.1% SDS, pH 8.3). Gels were incubated in transfer buffer (25 mM Tris, 192 mM glycine, pH 8.3, 20% methanol) containing 1mM EDTA for 10min, followed by incubation in transfer buffer for an additional 20min. Gels were then transferred (100V, 60m) to PVDF membrane, blocked in TBS (20mM Tris, 150mM NaCl, pH 7.5)/5% milk and probed with anti-Flag M2 antibody (Sigma) in TBST (20mM Tris, 150mM NaCl, 0.05% Tween20, pH 7.5)/5% milk. Membranes were washed in TBST and incubated with IRDye 680RD goat anti-mouse antibody (LI-COR) and imaged on a LI-COR Odyssey system. Intensity of individual bands were generated using ImageStudioLite or EmpiriaStudio (LI-COR). Fraction PilG-P was calculated from the intensity of the PilG-P band/total intensity of PilG bands (sum of both PilG bands).

### Statistical tests

To test significance, one-way ANOVA and Tukey’s post hoc tests were done where required using Python (version 3.8.5). Different groups values were considered significantly different with a p-value below 0.05.

## Acknowledgements

ZAM, XP, LT and AP are supported by the SNSF Projects grant number 310030_189084. MJK is supported by the EMBO postdoctoral fellowship ALTF 495-2020. JNE, YI, RP, HM and are supported by an NIH R01 grant number AI129547 and by the Cystic Fibrosis Foundation (495008).

---

**Supplementary Figure 1:**
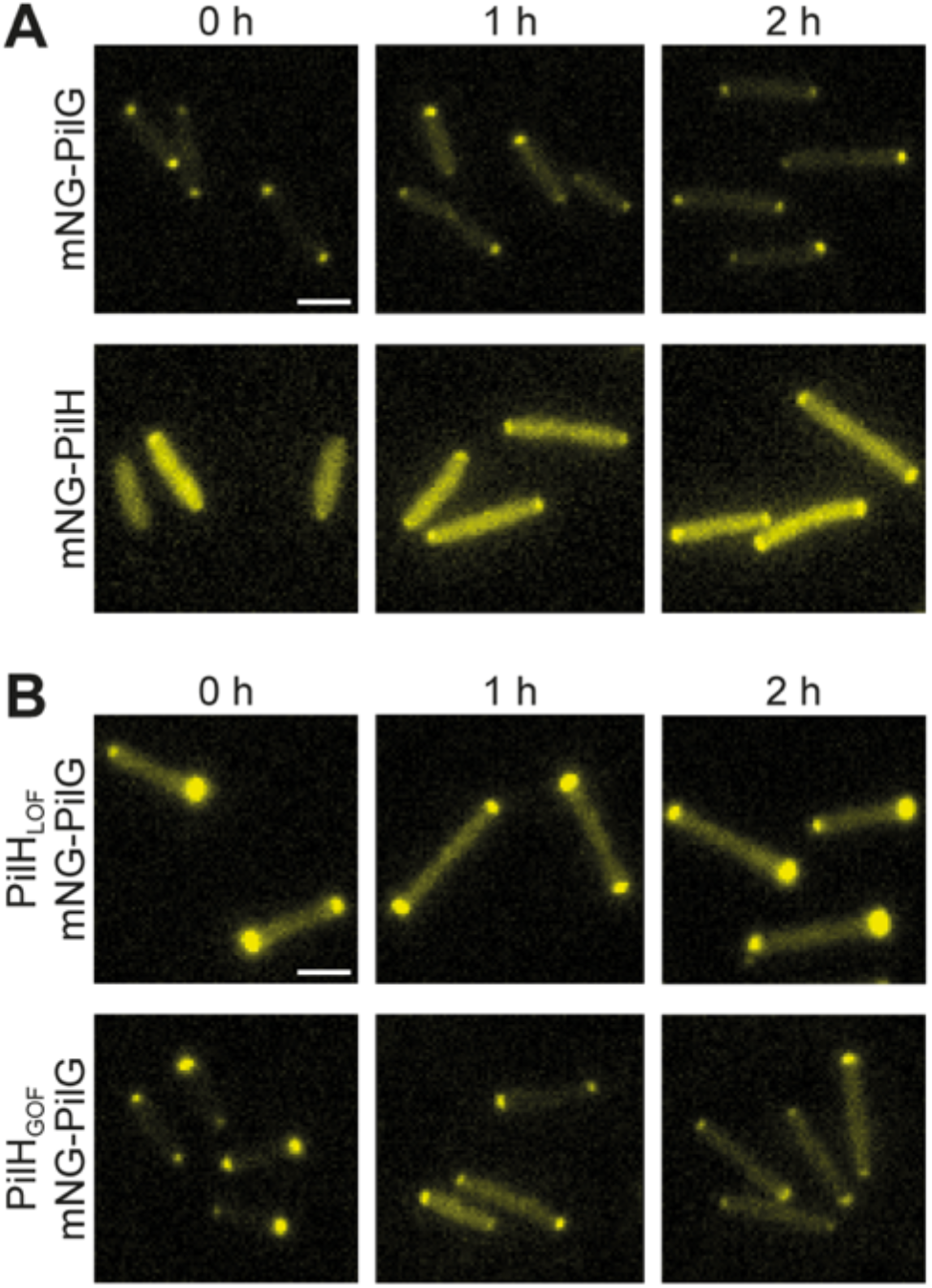
Example fluorescence microscopy images of mNG-PilG and mNG-PilH in cells grown on a surface. Cells were transferred from liquid culture to solid substrates and imaged immediately (0h) and after 1h and 2h in which the cells adapt to a surface-associated lifestyle. (A) Images correspond to data shown in Figure 1. (B) Images correspond to data shown in Figure 6. Scale bars, 2 μm.

**Supplementary Figure 2:**
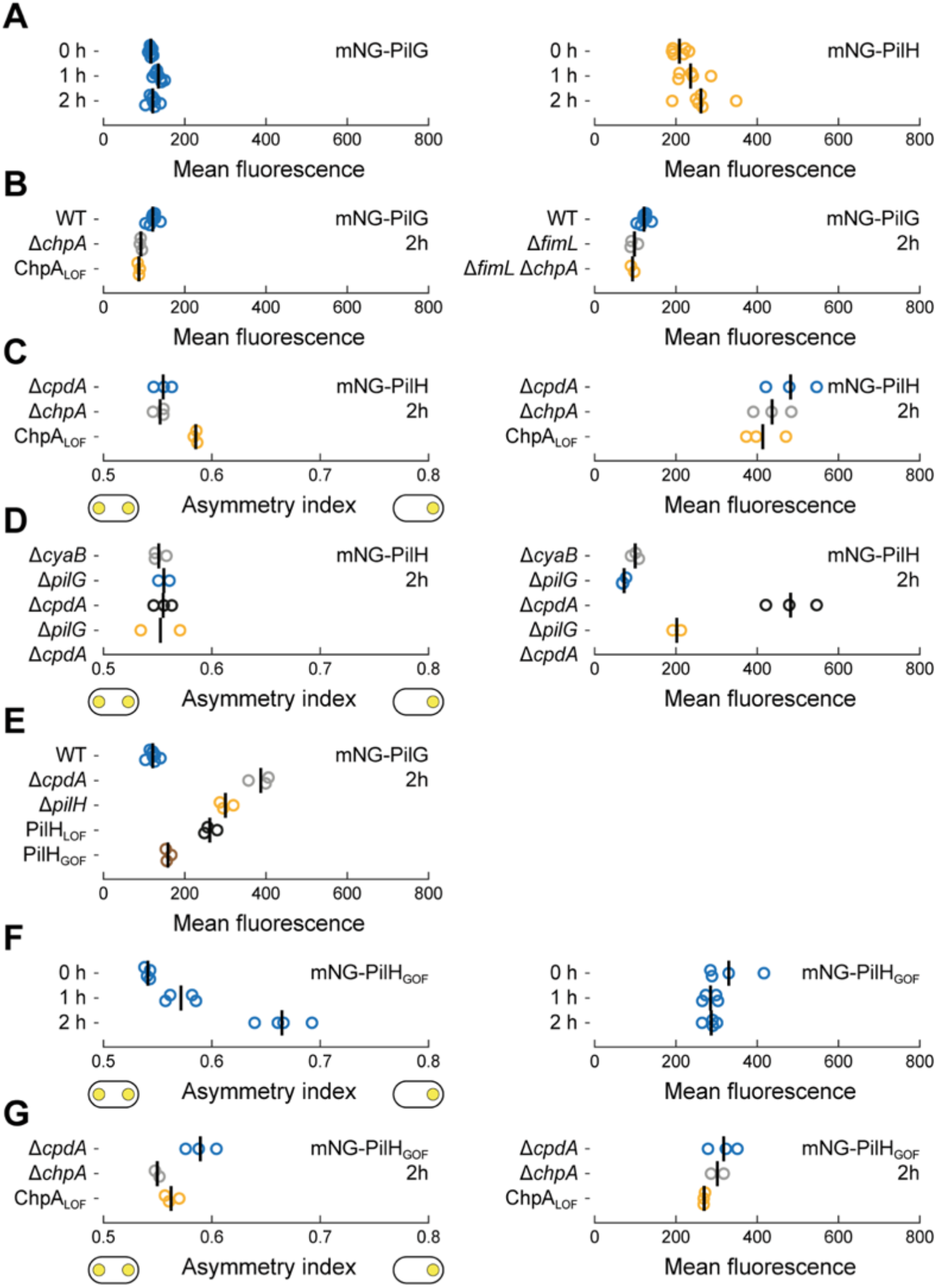
Mean cell fluorescence and asymmetry indices of mNG-PilG and mNG-PilH. Panels correspond to main figures as follows: (A) Figure 1 (B) Figure 2 (C) Figure 3 (D) Figure 5 (E) Figure 6 (F) Figure 7AB (G) Figure 7CD. Note, for panels C and G, *cpdA* was deleted in all display strains to rescue cAMP levels and mitigate the negative effects caused by low cAMP. Circles, median of each biological replicate. Vertical bars: mean across biological replicates.

**Supplementary Figure 3:**
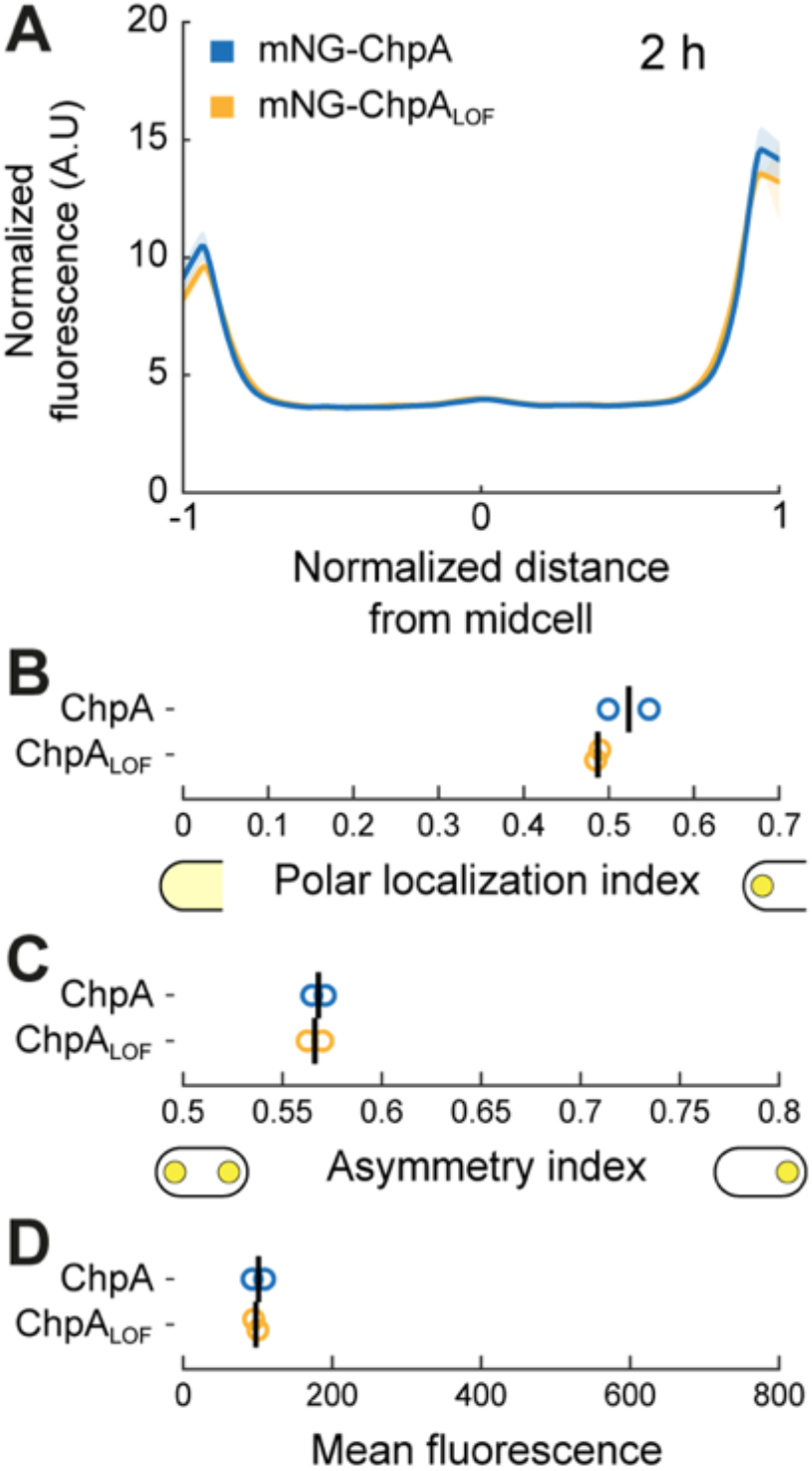
ChpA and ChpA_LOF_ similarly localize to the poles. (A) Average fluorescence profiles of mNG-ChpA after 2h surface growth. Note, wild-type mNG-tagged ChpA is non-functional (0 moving cells found in 97 tracked cells) like the loss-of-function mutant ChpA_LOF_ (2 moving cells found in 254 tracked cells). (B) Quantification of polar localization, (C) asymmetry index and (D) mean cellular fluorescence. Solid lines, mean normalized fluorescence profiles across biological replicates. Shaded area, standard deviation across biological replicates. Circles, median of each biological replicate. Vertical bars, mean across biological replicates.

**Supplementary Figure 4:**
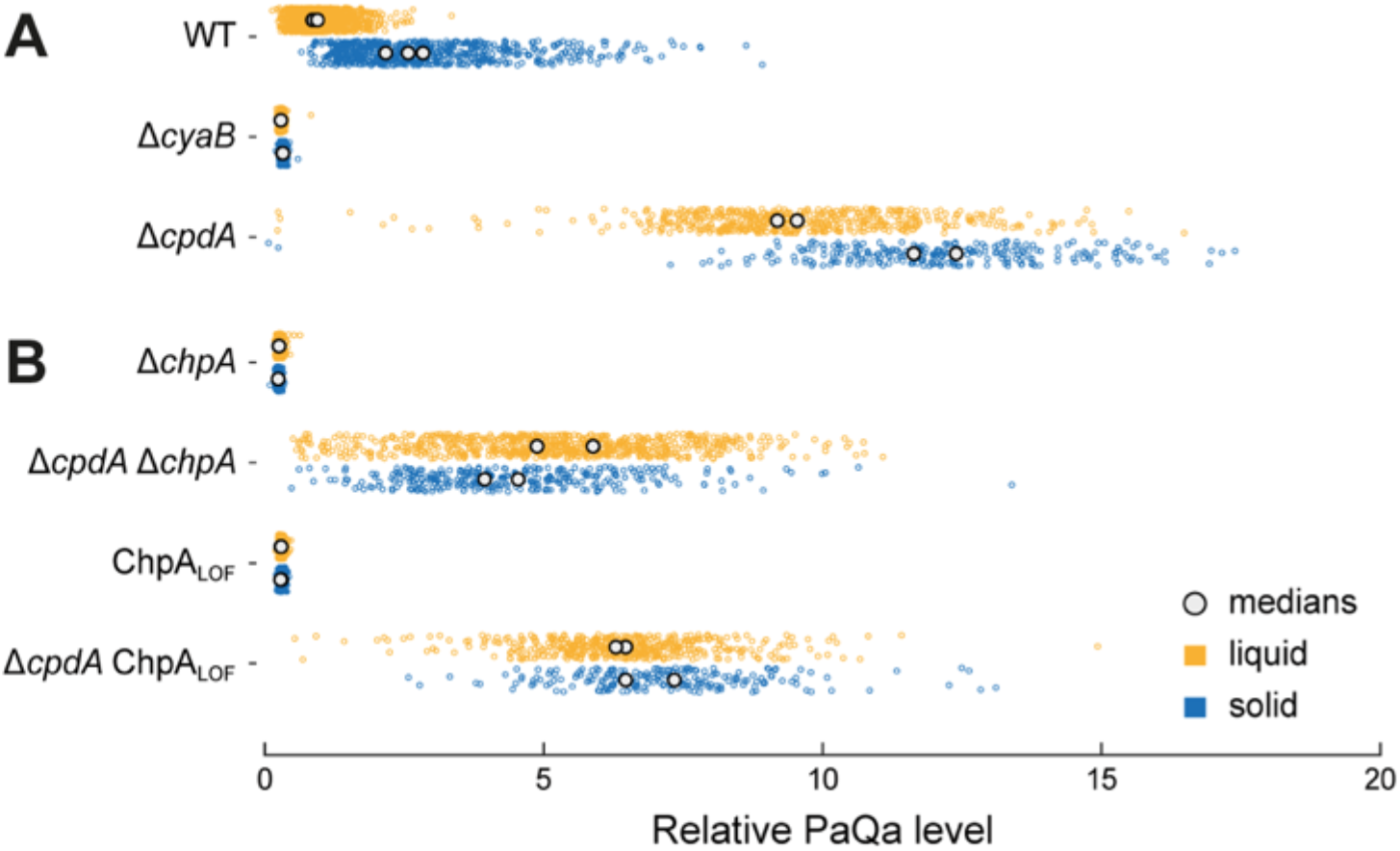
Quantification of cAMP levels of *chpA* mutants measured by PaQa-YFP reporter fluorescence. (A) Relative cAMP levels measured by PaQa reporter fluorescence in deletion mutants of the cAMP production cycle. Δ*cyaB* results in constitutively low and Δ*cpdA* in constitutively high cAMP levels. The cAMP level of Δ*cpdA* increases on surfaces, likely due to increased activity of the adenylate cyclase CyaB (11). (B) Relative cAMP levels of Δ*chpA*, the histidine kinase of the Chp system, and the ChpA loss-of-function mutant ChpA_LOF_. Note, deletion of *cpdA* partially rescues the low cAMP level of *chpA* mutants, however, cAMP levels don’t reach the high cAMP levels of Δ*cpdA*. Very likely, CyaB doesn’t get activated in *chpA cpdA* double mutants (11). Therefore, cAMP levels don’t increase upon surface contact. Colored circles represent measurements of single cells, white circles correspond to medians of biological replicates.

**Supplementary Figure 5:**
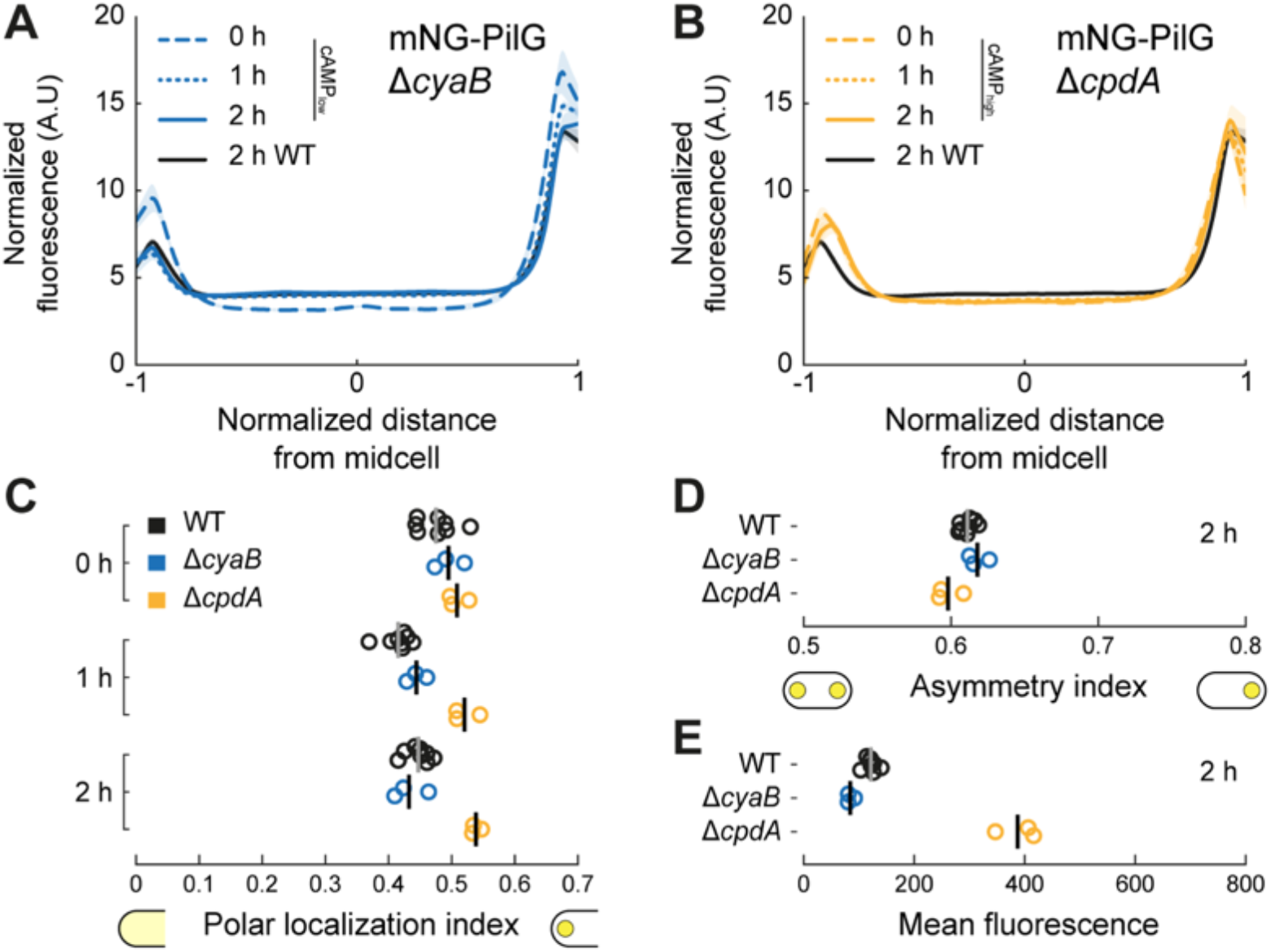
Time course of cAMP-dependent localization of mNG-PilG in cells grown on solid substrate. Average fluorescence profiles of mNG-PilG in (A) low and (B) high cAMP. Quantification of (C) polar localization, (D) asymmetry index and (E) mean cellular fluorescence. In low cAMP (Δ*cyaB*), the localization pattern as well as the decrease of polar localization over time is almost identical to WT. Data from Figure 1 included for comparison (black). Note, WT refers to a strain without *cyaB* or *cpdA* deletion. In high cAMP (Δ*cpdA*), polar localization of PilG is comparable to WT at 0h, however, remains high over time. The higher polar localization is due to increased localization to the dim pole (at x = −1, panel B) which is reflected by a slightly lower asymmetry index (D), i.e. more symmetric localization. The fluorescent signal of mNG-PilG, as a proxy for protein concentration, is largely unaffected by low cAMP, however, strongly increased in high cAMP. Solid lines, mean normalized fluorescence profiles across biological replicates. Shaded area, standard deviation across biological replicates. Circles, median of each biological replicate. Vertical bars, mean across biological replicates.

**Supplementary Figure 6:**
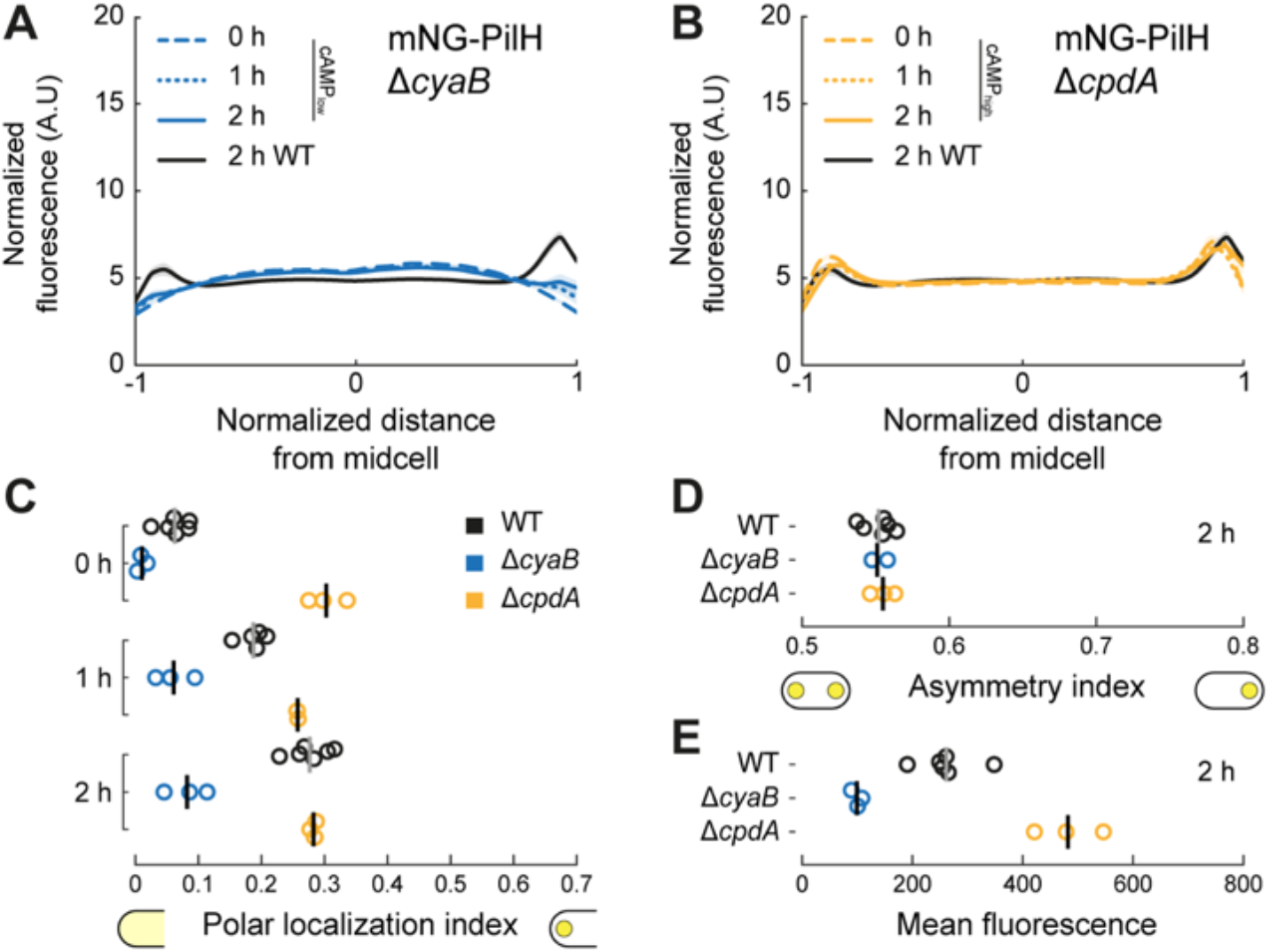
Time course of cAMP-dependent localization of mNG-PilH in cells grown on solid substrate. Average fluorescence profiles of mNG-PilH in (A) low and (B) high cAMP, respectively. Quantification of (C) polar localization, (D) asymmetry index and (E) mean cellular fluorescence. In low cAMP *(ΔcyaB)*, polar localization is generally reduced, however, recruitment of PilH to the poles over time still takes place. In high cAMP *(ΔcpdA)*, polar localization is always as high as in WT after 2h surface growth. Data from Figure 1 included for comparison (black). Note, WT refers to a strain without *cyaB* or *cpdA* deletion. The fluorescent signal of mNG-PilH, as a proxy for protein concentration, is proportional to the cAMP level in *cyaB* and *cpdA* mutants (cf. Supplementary Figure 4A). Solid lines, mean normalized fluorescence profiles across biological replicates. Shaded area, standard deviation across biological replicates. Circles, median of each biological replicate. Vertical bars, mean across biological replicates.

**Supplementary Figure 7:**
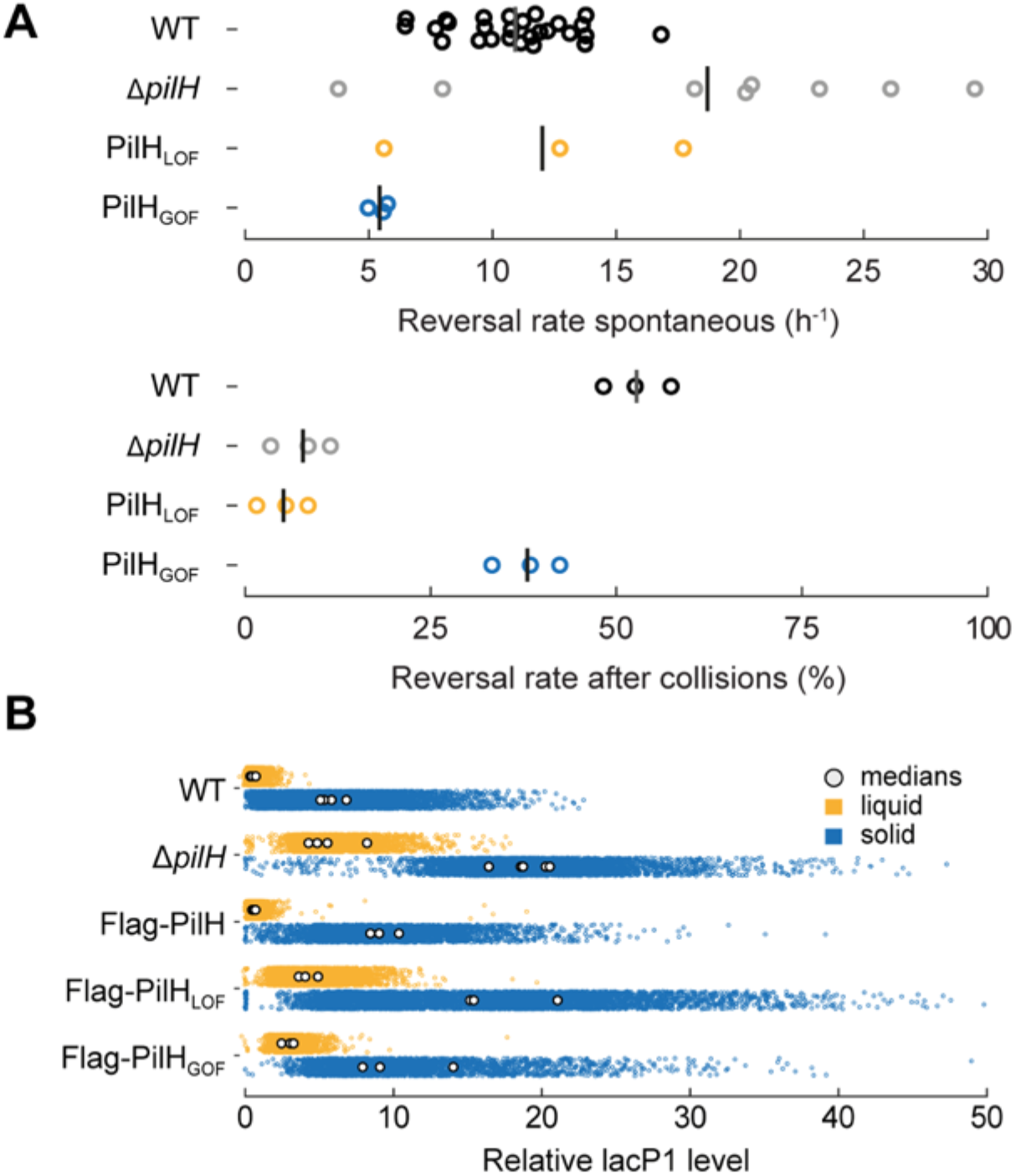
Reversal rates and cAMP production of loss- and gain-of-function mutants of PilH. (A) Spontaneous reversal rates and reversal rates after collision of PilH_LOF_ and PilH_GOF_ mutants. Δ*pilH* and PilH_LOF_ show low to almost no reversals (see also (9)). PilH_GOF_ rescues reversal rates, however, doesn’t fully restore wild-type rates. Because of high pili number in mutants with high cAMP level resulting in erratic movement of cells, spontaneous reversals rates are not measured very precisely by our algorithm. Manually counted reversal rates after collision give a better estimate. (B) Quantification of cAMP levels measured by PlacP1-YFP reporter fluorescence. cAMP levels of PilH_LOF_ are comparable to Δ*pilH*, whereas PilH_GOF_ partially rescues cAMP to intermediate levels between WT and Δ*pilH* or PilH_LOF_. Colored circles represent measurements of single cells, white circles correspond to medians of biological replicates.

**Supplementary Figure 8:**
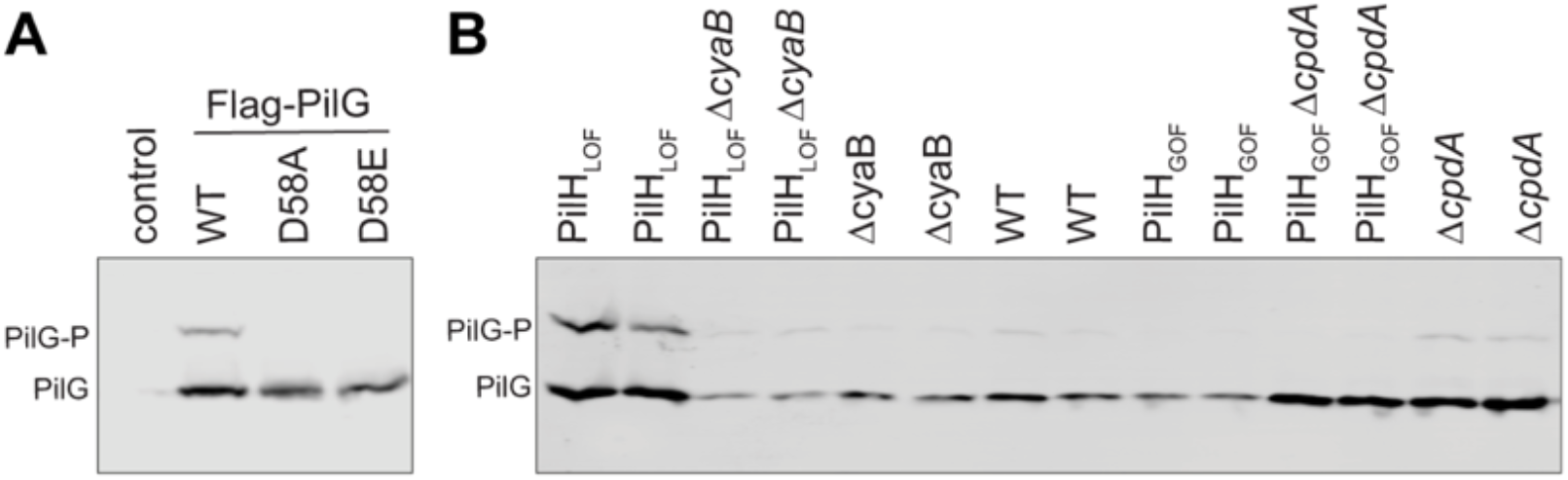
Example PhosTag™ western blots. (A) Representative gel of PilG and corresponding point mutants that can’t get phosphorylated. A slower migrating band corresponding to PilG-P is detected in whole cell lysates from wild type cells expressing Flag-PilG but not in cells expressing PilG with mutations in the phospho-accepting site D58. Control, wild-type PilG without Flag tag. (B) Representative gel of PilG in PilH point mutants and mutants with altered cAMP levels (Δ*cpdA*, Δ*cyaB*), corresponding to data shown in Figure 7EF.

**Supplementary Figure 9:**
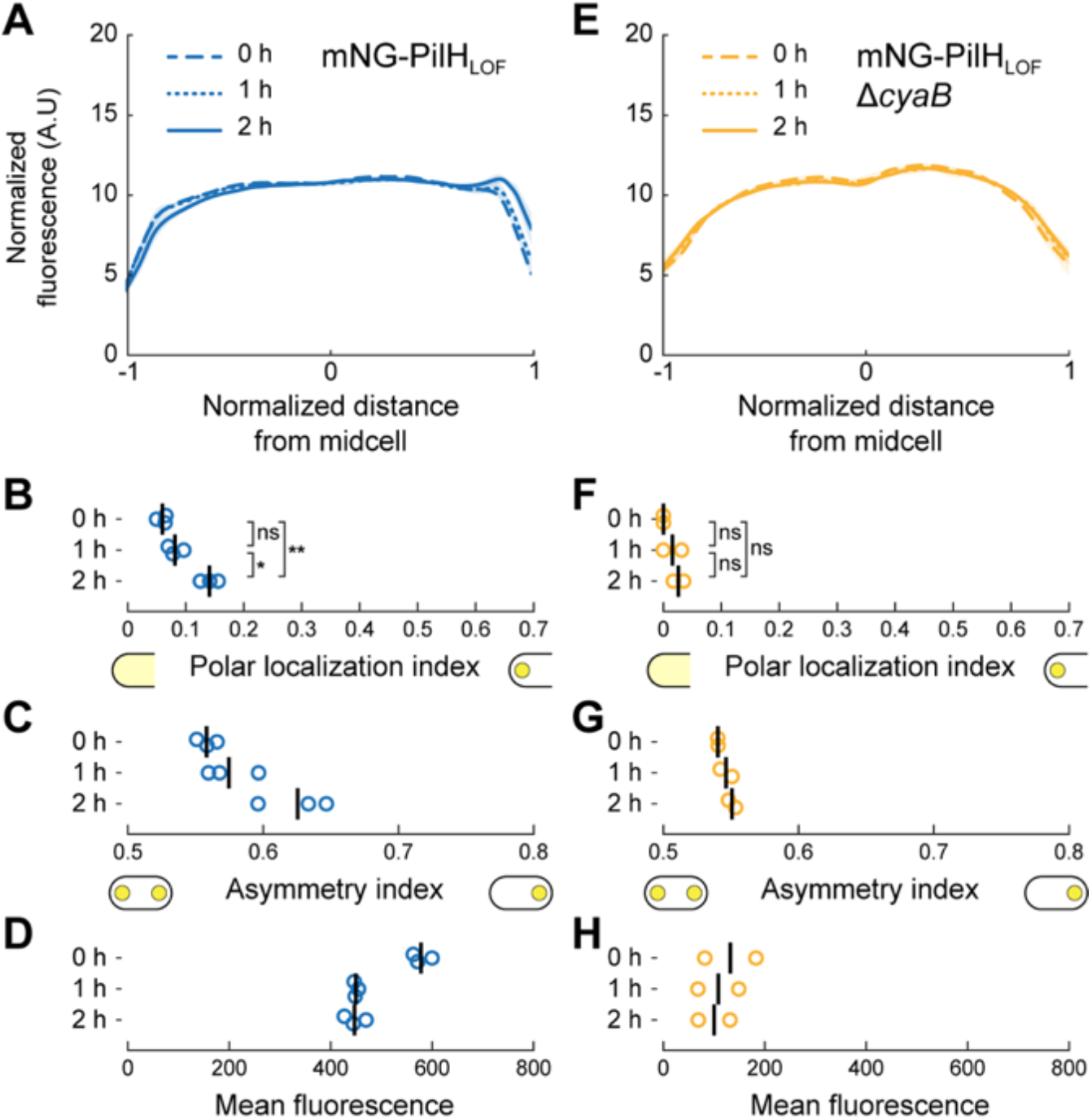
PilH locked in its inactive conformation is recruited to the poles upon surface contact. (A) Time course of localization profiles of mNG-PilH_LOF_ in cells grown on solid substrate. Quantification of corresponding (B) polar localization, (C) asymmetry index and (D) mean cellular fluorescence. Like PilH_wt_ and PilH_GOF_, PilH_LOF_ gets recruited to the poles over time (B). However, the effect is significantly less pronounced, possibly due to saturation effects because of high fluorescent signal. Note, loss-of-function mutation of *pilH* results in high cAMP like in Δ*pilH* (Supplementary Figure 7). (E-H) Same analysis in low cAMP (Δ*cyaB*). Polar recruitment may take place but the effect is too weak to be measured clearly (F). Solid lines, mean normalized fluorescence profiles across biological replicates. Shaded area, standard deviation across biological replicates. Circles, median of each biological replicate. Vertical bars, mean across biological replicates. *, p<0.05; **, p≤0.001; ns, not significant.

## Movies

**Supplementary movie 1: Single twitching *Pseudomonas aeruginosa* cell with fluorescently labelled type IV pili.** During the reversal of twitching direction T4P disappear at the initially leading cell pole (upper pole) and reappear at the opposite cell pole which becomes the new leading cell pole. Timestamp, min:sec; scale bar, 2 μm.

