## Supplement Table 1 for "*Pseudomonas aeruginosa* mechanosensing controls cell polarity during twitching by activating two antagonistic response regulators"

### Supporting Information

**Supplementary Table 1:** Strains used in this study. LOF, loss-of-function mutant. GOF, gain-of-function mutant.

| **Name and relevant genotype** | **Source / Reference** | **Identifier** |
|---|---|---|
| Pseudomonas aeruginosa PAO1 | (1) | ATCC 15692 |
| Escherichia coli DH5α (hsdR rec lacZYA ϕ80 lacZM15) | Invitrogen | Na |
| Escherichia coli strain S17.1 (thi pro hsdR recA RP4-2(Tc::Mu)(Km::Tn7)) | Stratagene | Na |
| PAO1 Δ*fliC* (in-frame deletion of PA1092) | (2) | 177 |
| PAO1 Δ*pilG* (in-frame deletion of PA0408) | (2) | 226 |
| PAO1 Δ*pilH* (in-frame deletion of PA0409) | (3) | 178 |
| PAO1 Δ*chpA* (in-frame deletion PA0413) | (1) | 170 |
| PAO1 Δ*cyaB* (in-frame deletion of PA3217) | (4) | 174 |
| PAO1 Δ*cpdA* (in-frame deletion of PA4969) | (4) | 180 |
| PAO1 Δ*fimL* (in-frame deletion of PA1822) | (5) | 179 |
| PAO1 Δ*fliC* Δ*cyaB* | (6) | 326 |
| PAO1 Δ*fliC* Δ*cpdA* | (6) | 337 |
| PAO1 Δ*fliC* Δ*pilG* | (6) | 330 |
| PAO1 Δ*fliC* Δ*pilH* | (6) | 232 |
| PAO1 Δ*fliC* Δ*pilG* Δ*cpdA* | (6) | 459 |
| PAO1 Δ*fliC* Δ*pilG* Δ*pilH* | this study | 553 |
| PAO1 Δ*fliC* Δ*pilG* Δ*pilH* Δ*cpdA* | this study | 1278 |
| PAO1 Δ*fliC* Δ*chpA* Δ*cpdA* | this study | 342 |
| PAO1 Δ*fliC* $ChpA_{ΔHK}$ (deletion of the histidine kinase domain, residues 1943-2176, parent for $ChpA_{LOF}$) | this study | 1511 |
| PAO1 Δ*fliC* ΔcpdA $ChpA_{ΔHK}$ | this study | 1512 |
| PAO1 Δ*fliC* $ChpA_{LOF}$ (loss-of-function mutations D2086A, D2087A, G2088A) | This study, based on (2) | 1536 |
| PAO1 Δ*fliC* Δ*cpdA* $ChpA_{LOF}$ | this study | 1537 |
| PAO1 Δ*fliC* $PilH_{LOF}$ (loss-of-function mutation D52A) | this study | 1172 |
| PAO1 Δ*fliC* $PilH_{LOF}$ Δ*cyaB* | this study | 1166 |
| PAO1 Δ*fliC* $PilH_{GOF}$ (loss-of-function mutation D52E) | this study | 1155 |
| PAO1 Δ*fliC* $PilH_{GOF}$ Δ*cyaB* | this study | 1167 |
| PAO1 Δ*fliC* mNG-PilG (N-terminal fluorescent fusion to mNeonGreen, GGGGG linker, native locus) | (6) | 923 |
| PAO1 Δ*fliC* mNG-PilG Δ*cyaB* | this study | 1098 |
| PAO1 Δ*fliC* mNG-PilG Δ*cpdA* | this study | 1429 |

| | | |
|---|---|---|
| PAO1 Δ*fliC* mNG-PilH (N-terminal fluorescent fusion to mNeonGreen, GGGGG linker, native locus) | (6) | 315 |
| PAO1 Δ*fliC* mNG-PilH Δ*cyaB* | this study | 1095 |
| PAO1 Δ*fliC* mNG-PilH Δ*cpdA* | this study | 1055 |
| PAO1 Δ*fliC* mNG-ChpA (N-terminal fluorescent fusion to mNeonGreen, GGGGG linker, native locus) | this study | 312 |
| PAO1 Δ*fliC* mNG-$ChpA_{ΔHK}$ | this study | 1510 |
| PAO1 Δ*fliC* mNG-$ChpA_{LOF}$ | this study | 1535 |
| PAO1 Δ*fliC* mNG-PilG Δ*chpA* | this study | 1423 |
| PAO1 Δ*fliC* mNG-PilG Δ*chpA* Δ*cpdA* | this study | 1464 |
| PAO1 Δ*fliC* mNG-PilG $ChpA_{ΔHK}$ | this study | 1517 |
| PAO1 Δ*fliC* mNG-PilG $ChpA_{LOF}$ | this study | 1542 |
| PAO1 Δ*fliC* mNG-PilG $ChpA_{ΔHK}$ Δ*cpdA* | this study | 1518 |
| PAO1 Δ*fliC* mNG-PilG $ChpA_{LOF}$ Δ*cpdA* | this study | 1578 |
| PAO1 Δ*fliC* mNG-PilG Δ*fimL* | this study | 1045 |
| PAO1 Δ*fliC* mNG-PilG Δ*fimL* Δ*chpA* | this study | 1433 |
| PAO1 Δ*fliC* mNG-PilG Δ*pilH* | this study | 969 |
| PAO1 Δ*fliC* mNG-PilG $PilH_{LOF}$ | this study | 1023 |
| PAO1 Δ*fliC* mNG-PilG $PilH_{GOF}$ | this study | 1404 |
| PAO1 Δ*fliC* mNG-PilH Δ*chpA* | this study | 1058 |
| PAO1 Δ*fliC* mNG-PilH Δ*chpA* Δ*cpdA* | this study | 1406 |
| PAO1 Δ*fliC* mNG-PilH $ChpA_{LOF}$ | this study | 1582 |
| PAO1 Δ*fliC* mNG-PilH $ChpA_{LOF}$ Δ*cpdA* | this study | 1539 |
| PAO1 Δ*fliC* mNG-PilH Δ*pilG* | this study | 916 |
| PAO1 Δ*fliC* mNG-PilH Δ*pilG* Δ*cpdA* | this study | 1041 |
| PAO1 mNG-$PilH_{LOF}$ | this study | 990 |
| PAO1 mNG-$PilH_{LOF}$ Δ*cyaB* | this study | 1360 |
| PAO1 mNG-$PilH_{GOF}$ | this study | 989 |
| PAO1 Δ*fliC* mNG-$PilH_{GOF}$ | this study | 1355 |
| PAO1 Δ*fliC* mNG-$PilH_{GOF}$ Δ*cpdA* | this study | 1467 |
| PAO1 Δ*fliC* mNG-$PilH_{GOF}$ Δ*cpdA* Δ*chpA* | this study | 1485 |
| PAO1 Δ*fliC* mNG-$PilH_{GOF}$ $ChpA_{ΔHK}$ | this study | 1515 |
| PAO1 Δ*fliC* mNG-$PilH_{GOF}$ $ChpA_{ΔHK}$ Δ*cpdA* | this study | 1516 |
| PAO1 Δ*fliC* mNG-$PilH_{GOF}$ $ChpA_{LOF}$ | this study | 1540 |
| PAO1 Δ*fliC* mNG-$PilH_{GOF}$ $ChpA_{LOF}$ Δ*cpdA* | this study | 1541 |
| PAO1 Δ*fliC* PaQa | (6) | 764 |
| PAO1 Δ*fliC* Δ*cpdA* PaQa | (6) | 867 |
| PAO1 Δ*fliC* Δ*cyaB* PaQa | (6) | 866 |
| PAO1 Δ*fliC* Δ*chpA* PaQa | this study | 1590 |

| | | |
|---|---|---|
| PAO1 Δ*fliC* Δ*chpA* Δ*cpdA* PaQa | this study | 1591 |
| PAO1 Δ*fliC* ChpA$_{LOF}$ PaQa | this study | 1580 |
| PAO1 Δ*fliC* ChpA$_{LOF}$ Δ*cpdA* PaQa | this study | 1581 |
| PAO1 Δ*fliC* PilA$_{A86C}$ (cysteine substitution for maleimide labelling, single point mutation of chromosomal PilA) | this study | 1210 |
| PAO1 3xFlag-PilG | this study | HM540 |
| PAO1 3xFlag-PilG$_{D58A}$ (non-phosphorylatable mutant) | this study | HM543 |
| PAO1 3xFlag-PilG$_{D58E}$ (non-phosphorylatable mutant) | this study | HM545 |
| PAO1 3xFlag-PilG Δ*cpdA* | this study | YI974 |
| PAO1 3xFlag-PilG Δ*cyaB* | this study | HM620 |
| PAO1 3xFlag-PilG Δ*pilH* | this study | HM587 |
| PAO1 3xFlag-PilG PilH$_{LOF}$ | this study | HM590 |
| PAO1 3xFlag-PilG PilH$_{GOF}$ | this study | HM592 |
| PAO1 3xFlag-PilG PilH$_{GOF}$ Δ*cpdA* | this study | HM664 |
| PAO1 PlacP1-YFP POXB20-mKate2 | this study | HM413 |
| PAO1 3xFlag-PilH PlacP1-YFP POXB20-mKate2 | this study | HM438 |
| PAO1 3xFlag-PilH$_{LOF}$ PlacP1-YFP POXB20-mKate2 | this study | HM439 |
| PAO1 3xFlag-PilH$_{GOF}$ PlacP1-YFP POXB20-mKate2 | this study | HM440 |
| PAO1 Δ*pilH* PlacP1-YFP POXB20-mKate2 | this study | HM422 |
| PAO1 Δ*cpdA* PlacP1-YFP POXB20-mKate2 | this study | HM419 |
| PAO1 Δ*cyaB* PlacP1-YFP POXB20-mKate2 | this study | HM416 |
| PAO1 3xFlag-PilG Δ*pilH* PlacP1-YFP POXB20-mKate2 | this study | HM603 |
| PAO1 3xFlag-PilG Δ*cpdA* PlacP1-YFP POXB20-mKate2 | this study | HM605 |
| PAO1 3xFlag-PilG Δ*cyaB* PlacP1-YFP POXB20-mKate2 | this study | HM628 |
| PAO1 3xFlag-PilG PilH$_{LOF}$ PlacP1-YFP POXB20-mKate2 | this study | HM609 |
| PAO1 3xFlag-PilG PilH$_{GOF}$ PlacP1-YFP POXB20-mKate2 | this study | HM611 |
| PAO1 3xFlag-PilG PilH$_{GOF}$ Δ*cpdA* PlacP1-YFP POXB20-mKate2 | this study | HM678 |

**Supplementary Table 2:** Plasmids used in this study. All displayed pEX18 and pEx100T vectors are suicide vectors for marker-free in-frame deletion or insertion. Deletions are marked by a Δ in front of the gene name.

| **Name and relevant information** | **Source / Reference** | **Identifier** |
|---|---|---|
| pEX100TAP (Suicide vector based on pUC19, Amp$^R$, ColE1 ori (*E. coli*), *oriT*, *sacB*, *lacZα*) | (7) | Na |
| pEX18AP (Suicide vector based on pUC18, Amp$^R$, ColE1 ori (*E. coli*), *oriT*, *sacB*, *lacZα*) | (8) | Na |
| pEX18GM (Suicide vector based on pUC18, Gm$^R$, ColE1 ori (*E. coli*), *oriT*, *sacB*, *lacZα*) | (8) | Na |

| | | |
|---|---|---|
| pUCP18-PaQa (fluorescent reporter for cAMP level: YFP controlled by *PaQa* promoter (PA1867 and PA1868) and mKate2 controlled by *rpoD* promoter (PA0576) as reference. | (9) | pAP02.2 |
| pUC18_PlacP1-YFP/POXB20-mKate2 (fluorescent reporter for cAMP level: YFP controlled by the synthetic *LacP1* promoter and mKate2 controlled by POXB20 promoter (Oxford Genetics Ltd. (UK), Sigma) as reference. | this study | YI996 |
| pEx100TAP-Δ*fliC* (PA1092) | (2) | pJB215 |
| pEx100TAP-Δ*cpdA* (PA4969) | (10) | pJTW033 |
| pEX18GM-Δ*cpdA* (PA4969) | (6) | pMK019 |
| pEx100TAP-Δ*cyaB* (PA3217) | (10) | pJTW031 |
| pEX18GM-Δ*cyaB* (PA3217) | (6) | pMK018 |
| pEx100TAP-Δ*pilG* (PA0408) | (2) | PJB118 |
| pEx100TAP-Δ*pilH* (PA0409) | (2) | PJB119 |
| pEX18AP-Δ*pilGH* (PA0408 and PA0409 including intergenic region) | this study | pXP322 |
| pEX18GM-*chpA*$_{\Delta HK}$ (deletion of the histidine kinase domain of PA0413, residues 1943-2176) | this study | pMK056 |
| pEX18GM-*chpA*$_{LOF}$ (insertion of the histidine kinase domain of PA0413 with substituted residues D2086A, D2087A, G2088A) | this study, based on (2) | pMK057 |
| pEX18AP-*pilH*$_{LOF}$ (PA0409 with substituted residue D52A) | this study | pMK012 |
| pEX18AP-*pilH*$_{GOF}$ (PA0409 with substituted residue D52E) | this study | pMK013 |
| pEx100TAP-mNG-PilH (PA0409 N-terminus fused with mNeonGreen, GGGGG linker) | (6) | pXP125 |
| pEX18GM-mNG-PilG (PA0408 N-terminus fused with mNeonGreen, GGGGG linker) | (6) | YI883 |
| pEx100TAP-mNG-ChpA (PA0413 N-terminus fused with mNeonGreen, GGGGG linker) | this study | pXP117 |
| pEX18GM-PilA-A86C (cysteine-labelled PA4525) | this study | pMK025 |
| pEX18GM-mNG-PilH$_{LOF}$ (PA0409 with substituted residue D58A N-terminus fused with mNeonGreen, GGGGG linker) | this study | YI926 |
| pEx100TAP-mNG-PilH$_{GOF}$ (PA0409 with substituted residue D58E N-terminus fused with mNeonGreen, GGGGG linker) | this study | HM83 |
| pEX18GM-3xFlag-PilG (PA0408 N-terminus fused with 3xFlag, GGGGG linker) | this study | HM524 |
| pEX18GM-3xFlag-PilG$_{D58A}$ (PA0408 D58A non-phosphorylatable mutation, N-terminus fused with 3xFlag, GGGGG linker) | this study | HM526 |
| pEX18GM-3xFlag-PilG$_{D58E}$ (PA0408 D58E non-phosphorylatable mutation, N-terminus fused with 3xFlag, GGGGG linker) | this study | HM536 |
| pEx100TAP-3xFlag-PilH (PA0409 N-terminus fused with 3xFlag, GGGGG linker) | this study | HM118 |
| pEx100TAP-3xFlag-PilH$_{LOF}$ (PA0409 with substituted residue D52A N-terminus fused with 3xFlag, GGGGG linker) | this study | HM169 |

| | | |
|---|---|---|
| pEx100TAP-3xFlag-PilH$_{GOF}$ (PA0409 with substituted residue D52E N-terminus fused with 3xFlag, GGGGG linker) | this study | HM171 |
| pEX18GM-3xFlag-PilG Δ*pilH* (PA0408 N-terminus fused with 3xFlag, GGGGG linker, and in-frame deletion of PA0409) | this study | HM577 |
| pEX18GM 3xFlag-PilG PilH$_{LOF}$ (PA0408 N-terminus fused with 3xFlag, GGGGG linker, PA0409 with substituted residue D52A) | this study | HM583 |
| pEX18GM 3xFlag-PilG PilH$_{GOF}$ (PA0408 N-terminus fused with 3xFlag, GGGGG linker, PA0409 with substituted residue D52E) | this study | HM581 |

**Supplementary Table 3:** Oligonucleotides used in this study.

| **Identifier** | **Sequence** | **Purpose** |
|---|---|---|
| oXP794 | ATG ACC ATG ATT ACG AAT TCC AGT TCG TGC AGC GG | Generation of pXP322 |
| oXP795 | GCT GCG ACG GGC TCA CAT GTT CGC CCT ATA TCG AC | Generation of pXP322 |
| oXP796 | TAT AGG GCG AAC ATG TGA GCC CGT CGC AGC | Generation of pXP322 |
| oXP797 | GCC TGC AGG TCG ACT CTA GAA ATG AAG GGT TGC AGT GC | Generation of pXP322 |
| oMK136 | TGC ATG CCT GCA GGT CGA CTG ATC CTG CAC ACC CTC AAG G | Generation of pMK056/57 |
| oMK137 | GGT TCA CCG ACT GCA ACT GCG AAT AGC GG | Generation of pMK056 |
| oMK138 | GCA GTT GCA GTC GGT GAA CCG GGC GCT G | Generation of pMK056 |
| oMK139 | CAG CTA TGA CCA TGA TTA CGA ACC GTC CAT GCG CGG CAT C | Generation of pMK056/57 |
| oMK142 | CCG GCC GCG GCC GCG GAG AGG GTG AGG AGG ATG | Generation of pMK057 |
| oMK143 | CTC TCC GCG GCC GCG GCC GGC ATC CGC CTC GAC | Generation of pMK057 |
| oMK140 | TGC TGG AGA ACC TCG AAC TG | Check c*hpA*$_{HK}$ locus |
| oMK141 | TGA TCA TGA TGA TCG GCA GG | Check c*hpA*$_{HK}$ locus |
| oMK039 | CAT GCC TGC AGG TCG ACT CAC AGA GGG ATG ACC CGG | Generation of pMK052 |
| oMK042 | GCT ATG ACC ATG ATT ACG CGG TGG AAG TGG AAG TGG | Generation of pMK052 |
| oMK062 | GAG CCG GAT TGC AAC AAG TTG GGT GTA ATT GC | Generation of pMK052 |
| oMK063 | AAC TTG TTG CAA TCC GGC TCG ACG CCG | Generation of pMK052 |
| oLT040 | GTA TCG ACC GGG CAA TTG C | Check *pilA* locus |
| oLT043 | CTC TTG GGT GGA CTT GTC | Check *pilA* locus |
| YIp276 | GGC GCG GCA TCA TGA TGG CGA CGA AAA TGA TGT TC | Generation of PilG$_{D58A}$ mutation |
| YIp277 | GAA CAT CAT TTT CGT CGC CAT CAT GAT GCC GCG CC | Generation of PilG$_{D58A}$ mutation |

| | | |
|---|---|---|
| YIp284 | AGG CGC GGC ATC ATG ATT TCG ACG AAA ATG ATG TT | Generation of $PilG_{D58E}$ mutation |
| YIp285 | AAC ATC ATT TTC GTC GAA ATC ATG ATG CCG CGC CT | Generation of $PilG_{D58E}$ mutation |
| YIp280 | CGG GCA TGA CGA TGG CCA TCA GGA CCA CG | Generation of $PilH_{D52A}$ mutation |
| YIp281 | CGT GGT CCT GAT GGC CAT CGT CAT GCC CG | Generation of $PilH_{D52A}$ mutation |
| YIp282 | CCG GGC ATG ACG ATT TCC ATC AGG ACC AC | Generation of $PilH_{D52E}$ mutation |
| YIp283 | GTG GTC CTG ATG GAA ATC GTC ATG CCC GG | Generation of $PilH_{D52E}$ mutation |
| hm66 | GTA GTC ATC GAT TTT GTC ATC GTC TTT GTA GTC GGC GGC TTT GTC ATC GTC TTT GTA GTC GTT CGC CCT ATA TCG ACT | Generation of 3xFlag-PilG |
| hm67 | TAC AAA GAC GAT GAC AAA ATC GAT GAC TAC AAA GAC GAT GAC AAA GGC GGC GGC GGC GGC ATG GAA CAG CAA TCC GAC | Generation of 3xFlag-PilG |
| hm75 | TAT TTC GTG ATG GGG ATC CCA TGG CTC GTA TAA GCT TCA CCA CCA AGG ACC AG | Generation of 3xFlag-PilG ΔpilH |
| hm76 | CGA CGG CCA GTG CCA AGC TTT CGG GGC TGG GCG GCA GG | Generation of 3xFlag-PilG ΔpilH |
| hm52 | AAG CTT GGC ACT GGC CGTC | Generation of 3xFlag-PilG ΔpilH |
| hm53 | GGG ATC CCC ATC ACG AAA TAA G | Generation of 3xFlag-PilG ΔpilH |
